# Intrinsic Ionic Mechanisms Underlying Burst Firing in Spontaneously Active Dorsal Horn Parvalbumin Interneurons

**DOI:** 10.64898/2026.09.05.748938

**Authors:** Rian Fritz D. Jalandoni, Christopher Todrineau, Haoyi Qiu, Erik Cook, Arjun Krishnaswamy, Reza Sharif-Naeini, Anmar Khadra

**Affiliations:** Quantitative Life Sciences, McGill University, 550 Sherbrooke W., Montreal, H3A 1E3, QC, Canada; Department of Physiology, McGill University, 3655 Promenade Sir William Osler, Montreal, H3G 1Y6, QC, Canada

**Keywords:** Parvalbumin-expressing inhibitory interneurons, spontaneous firing activity, two-compartment Hodgkin–Huxley model, elliptic bursting, intrinsic vs circuit properties

## Abstract

Parvalbumin-expressing interneurons (PVINs) in the spinal dorsal horn play a key role in preventing touch inputs from engaging nociceptive pathways through fast inhibitory control. Although most PVINs are typically quiescent and require external input to fire, a subset of these neurons exhibits spontaneous activity, including isolated spikes and bursts. The mechanisms underlying this behavior in spontaneously active PVINs (spPVINs) remain unclear. To address this, we developed a stochastic two-compartment Hodgkin–Huxley (HH) type model, consisting of a soma and an axon initial segment (AIS), to investigate the effects of synaptic noise and intrinsic electrical properties of spPVINs in driving their spontaneous firing. The model incorporates two subthreshold currents: the M-type K^**+**^ current (***I***_***m***_) and the hyperpolarization-activated current (***I***_***h***_). The model revealed that, in the presence of Ornstein–Uhlenbeck noise, ***I***_***h***_ promotes spontaneous firing by enhancing noise-driven depolarizations. Model simulations closely reproduced the firing patterns of spPVINs observed experimentally, including irregular spiking and bursting. Bifurcation analysis showed that varying the applied current can lead to transitions between quiescent, tonic, and elliptic bursting regimes, allowing stochastic fluctuations to switch between these states, which gives rise to irregular spontaneous activity. Reducing the conductance of ***I***_***m***_ and increasing the conductance of the fast Na^**+**^ current (***I***_***Na***_) both enlarge the elliptic bursting regime, thereby enabling noise to promote bursting. Interestingly, when the model was incorporated into a circuit, the downstream inhibitory effects of spPVINs were enhanced by the expression of ***I***_***h***_. Taken together, these results demonstrate how intrinsic electrical properties of spPVINs shape their firing dynamics and circuit activity.

## Introduction

Neural circuits require the right balance between excitation and inhibition to process sensory information correctly. In pain processing, for example, defective inhibitory neurons can lead to chronic pain. Since ion channels control how neurons fire, any change in their behavior directly alters the spiking patterns that define a neuron’s role in the circuit.

In the spinal dorsal horn, parvalbumin-expressing inhibitory interneurons (PVINs) act as sentinels for incoming touch and pain signals [1, 2]. These fast-spiking neurons are mostly located at the border of laminae II and III, where they provide both presynaptic inhibition to sensory fibers and postsynaptic inhibition to local excitatory neurons [1–3]. This inhibition ensures that low-threshold mechanical inputs, like touch, do not activate polysynaptic pathways connected to pain projection neurons.

When PVINs are silenced in otherwise naive mice, the mice develop mechanical allodynia, or touch-evoked pain [1, 4]. On the other hand, activating PVINs in nerve-injured mice can alleviate their mechanical allodynia. These observations demonstrate that PVINs play a crucial role in preventing touch inputs from activating pain projection neurons, and that their dysfunction after nerve injury can lead to mechanical allodynia. Nerve injury causes this disinhibition through several distinct mechanisms. For example, after nerve injury, there is a decrease in the number of PVIN synapses onto their post-synaptic PKC*γ*-expressing neurons [1]. At the same time, retinoic acid receptor alpha (RAR*α*) signaling in PVINs has also been shown to affect the inhibitory synaptic output of these neurons [3]. Moreover, aside from synaptic changes, the intrinsic firing of PVINs also decreases [4, 5]. After injury, these neurons often lose their ability to fire at high, sustained frequencies, instead switching to an “adaptive” or burst-like pattern [2, 5]. This shift effectively removes the “gate” that prevents touch signals from being perceived as pain [1, 5].

PVINs are also responsible for setting the timing and synchrony of dorsal horn network activity. Nerve injury, moreover, decreases the recruitment of PVINs by low-threshold primary afferents [4, 6]. Thus, sensory processing requires PVINs to fire with high frequency and precision [4, 5], but how this occurs biophysically is not well understood.

Although PVINs rarely exhibit spontaneous firing in the absence of synaptic input, their diverse firing patterns under physiological conditions are important for regulating the strength and timing of inhibition in sensory circuits. PVINs exhibit tonic, adapting, or burst-like discharge, and these patterns are shaped by both intrinsic membrane conductances and synaptic network inputs. In particular, calcium-dependent potassium conductances, including SK channels, can strongly regulate PVIN firing mode and therefore their inhibitory output [7–10].

Spontaneously active PVINs (referred to hereafter as spPVINs) may be functionally important because they supply a baseline level of inhibition that helps maintain the sensory gate—the dorsal horn mechanism that blocks low-threshold touch signals from accessing pain pathways in the absence of input. An important unresolved question is whether spontaneous firing in these cells is driven primarily by extrinsic factors, such as ongoing synaptic input and network activity, or by intrinsic mechanisms associated with the expression and regulation of voltage-gated and calcium-dependent ion channels.

Conductance-based Hodgkin–Huxley (HH) type models provide important insights into the mechanisms underlying neuronal excitability by identifying the specific ionic conductances and biophysical processes responsible for experimentally observed electrophysiological behaviors [11, 12]. These models integrate both the intrinsic electrical properties of neurons, determined by the integration of the different ionic conductances expressed in these neurons, and the extrinsic influences arising from synaptic inputs and network interactions [13–16]. Furthermore, these conductance-based models can be formulated as multicompartmental representations that explicitly incorporate distinct anatomical regions of a neuron, including for example the dendrites, the soma, and the axon initial segment (AIS) [17–20]. This framework extends the scope of computational modeling by enabling investigations of how neuronal morphology, as well as distal and proximal synaptic activity, shape somatic membrane dynamics and action potential generation, where electrical activity is most commonly recorded experimentally.

To investigate the mechanisms underlying the spontaneous firing activity of spPVINs, we have developed in this study a modified two-compartment HH type model based on earlier work [8]. The two compartments represent the soma and AIS. This approach allowed us to simulate how specific ionic currents contribute to the fast-spiking behavior of PVINs. To account for ongoing synaptic bombardment, we incorporated an Ornstein– Uhlenbeck (OU) noise process into the model [21, 22], and compared it with intrinsic properties in terms of shaping the spontaneous activity of spPVINs. By applying bifurcation theory to our model, we then identified the various regimes of behavior and examined how variations in maximum conductances (such as those of the M-type K^+^ and fast Na^+^ currents) can induce transitions between them [5, 8]. Overall, this modeling study provides a mechanistic account of how these neurons operate under physiological conditions and how their dysfunction may contribute to circuit-level abnormalities observed in neuropathic pain.

## Methods

### Experimental recordings

#### Electrophysiology

Mice were anesthetized with an intraperitoneal injection of 2,2,2-Tribromoethanol (Avertin, 250 mg/kg) for deep anesthesia. The back of the mouse was shaved with an electric clipper, and a dorsal laminectomy was performed. A block containing the spinal cord was quickly transferred to a dissection dish perfused with an ice-cold (4^*°*^C) N-Methyl-D-Glucamine-based artificial cerebrospinal fluid (NMDG-ACSF) solution containing the following (in mM): 93 NMDG, 2.5 KCl, 1.25 NaH_2_PO_4_, 30 NaHCO_3_, 20 HEPES, 25 glucose, 2 thiourea, 5 Na-L-ascorbate, 3 Na-pyruvate, 12 N-acetyl-L-cysteine, 0.5 CaCl_2_ *·*2 H_2_O, and 10 MgSO_4_ *·*7 H_2_O (pH 7.3–7.4, adjusted with HCl 12 M), bubbled with 95% O_2_ and 5% CO_2_. The dura mater and pia were removed, and the spinal cord was secured onto an agar block using insect pins. Finally, a transverse slice (530–570 *µ*m) of the spinal cord was obtained using a vibratome (VT1200 S; Leica). Slices were transferred to a submerged chamber containing HEPES-based recovery ACSF for 20 min at 34^*°*^C, equilibrated with 95% O_2_ and 5% CO_2_, and were then maintained at room temperature until recording.

#### Targeted whole-cell patch clamp

Slices were transferred to a recording chamber and continuously superfused (2 mL/min) with oxygenated ACSF containing (in mM): 119 NaCl, 24 NaHCO_3_, 2.5 KCl, 1.25 NaH_2_PO_4_, 2 CaCl_2_, 2 MgCl_2_, and 12.5 glucose (bubbled with 95% O_2_ and 5% CO_2_; pH 7.3; 300*±*5 mOsm/kg measured). Patch pipettes were pulled from borosilicate glass capillaries (Harvard Apparatus) with a P-97 puller (Sutter Instruments). They were filled with a solution containing (in mM) 135 K-Gluconate, 6 NaCl, 2 MgCl_2_, 10 HEPES, 0.1 EGTA, 2 MgATP, 0.8 NaGTP (pH 7.3–7.4, adjusted with KOH; osmolarity, 300 mOsm, adjusted with sucrose) and had final tip resistances of 6–8 MΩ for whole-cell recording. Neurons were viewed with an upright microscope (Olympus) using a 40*×* water-immersion objective, infrared differential interference contrast (IR-DIC) and fluorescence. Recordings were made in whole-cell current clamp (holding potential at *−* 70 mV) from identified PVINs expressing tdTomato. Data were acquired with pClamp 10.0 software (Molecular Devices) using a MultiClamp 700B patch-clamp amplifier and a Digidata 1440A (Molecular Devices). Recordings were low-pass filtered online at 1 kHz, digitized at 20 kHz and stored on a PC using pClamp software (Molecular Devices). After obtaining the whole-cell recording configuration, access resistance and membrane capacitance were calculated based on the response to a 10 mV hyperpolarizing voltage step from a holding potential of *−* 70 mV. Pipette offset was zeroed, and the cell was excluded if a drift of more than 5 mV was noted.

Standardized current-clamp protocols were applied to each recorded cell. A gap-free 90-s-long recording with no injected current (*I*_*app*_ = 0 pA) was used to measure the resting membrane potential (mV). A 1-s-long step current injection from *−*200 to +200 pA (50 pA step increments) was used to measure the spike count and spiking duration.

#### Statistics

Experimental values are reported as mean *±* standard error of the mean (SEM), with *n* denoting the number of cells. Comparisons of passive membrane properties between spontaneously active and quiescent PVINs were performed using the two-sided Mann–Whitney *U* test, and differences were considered statistically significant at *p <* 0.05.

#### Distinct spontaneous activity phenotypes in PVINs

In the absence of an external current, we observed three distinct activity phenotypes across the dataset: quiescent, sparse-firing, and phasic-firing (Figure 1A). Among 25 naive PVINs, only eight exhibited sustained spontaneous activity (both sparse and phasic), suggesting that baseline excitability is tightly regulated and highly heterogeneous across the population. Quiescent PVINs (Figure 1A, top panel) are those that remain completely silent during the recording, whereas sparse-firing PVINs are those that exhibit spontaneous action potentials separated by relatively long interspike intervals (ISIs; Figure 1A, second and third panels). In contrast, phasic-firing PVINs are those that exhibit burst-like activity (Figure 1A, fourth and bottom panels) despite the absence of external current. For subsequent modeling and mechanistic analysis of these spontaneously active neurons (spPVINs), we selected a representative phasic-firing cell. The ISI histogram of this neuron reveals two distinct modes corresponding to intra-burst and inter-burst intervals (Figure 1B) [23]. Moreover, the L-shaped ISI return map (Figure 1C) further supports the presence of burst-firing dynamics and highlights the interplay between slow and fast processes underlying the transitions between quiescence, isolated spiking, and bursting. To test whether spontaneous firing reflected compromised recordings rather than genuine activity, we compared passive membrane properties between groups, classifying a cell as spontaneously active if it fired at least one spike during the 90-s gap-free recording. Spontaneously active cells were not depolarized at rest relative to quiescent cells (*−* 56.08 *±* 1.40 mV, *n* = 8, vs. *−* 55.45 *±* 1.76 mV, *n* = 17; Mann–Whitney *U, p* = 0.690), arguing against a leak-induced depolarization towards threshold. Input resistance was lower in spontaneously active cells (*R*_in_ = 243.72 *±* 35.41 MΩ, *n* = 8) than in quiescent cells (351.35 *±* 28.93 MΩ, *n* = 17), though this difference did not reach significance (*p* = 0.106) and our sample provides limited power to detect an effect of this magnitude. Spontaneous activity was therefore not attributable to recording artifact.

**Fig. 1.**
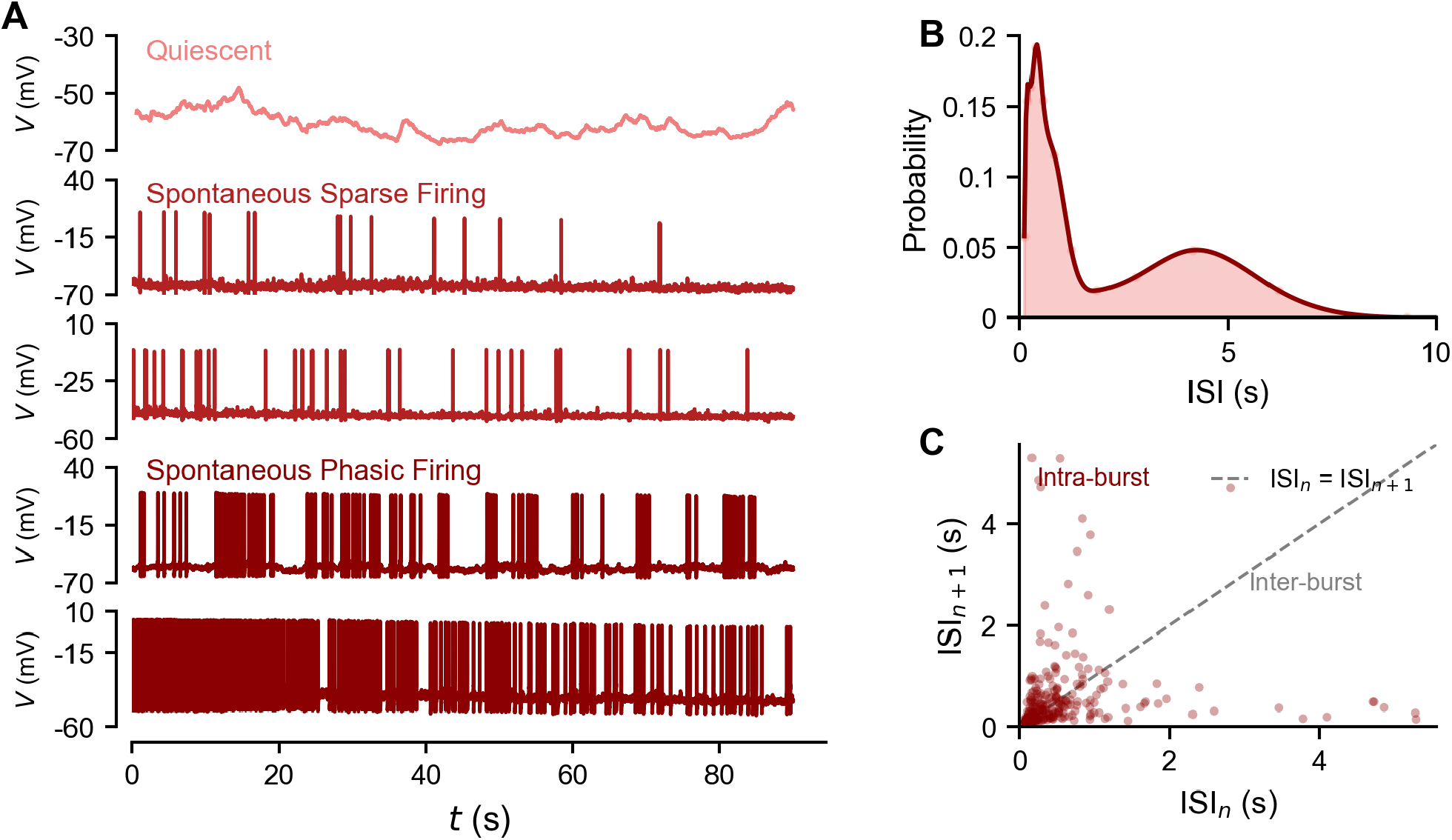
Spontaneous firing patterns and burst dynamics in spinal parvalbumin-expressing interneurons (PVINs). (A)Representative voltage recordings from PVINs in the absence of injected current. Three distinct activity phenotypes were observed: quiescent neurons (top panel) that remain near resting membrane potential, sparsely firing neurons (second and third panels) that produce isolated spikes at irregular intervals, and phasic neurons (fourth and bottom panels) that fire bursts of action potentials separated by silent periods. (B) Interspike interval (ISI) distribution of the spontaneously active phasic-firing PVIN shown in the fourth panel of A. The bimodal structure, with a sharp peak at short ISIs and a broader peak at longer ISIs, reflects two distinct timescales of firing: rapid intra-burst spiking and slower inter-burst intervals. (C) ISI return map (ISI_*n*+1_ vs. ISI_*n*_) from the same phasic-firing spPVIN shown in B, displaying an L-shaped profile that further supports the irregular, burst-dominated nature of this neuron.

### Mathematical model framework

#### Conductance-based two-compartment HH model

The computational framework developed here extends a previously established one-compartment Hodgkin– Huxley (HH) model [8] to more accurately capture the electrophysiological features of spPVINs. Two biophysical features were added: (1) a second compartment representing the axon initial segment (AIS; Figure 2A), and an Ornstein–Uhlenbeck (OU) noise process to represent stochastic synaptic bombardment and membrane fluctuations. The AIS included in this two-compartment HH model (Figure 2A) is the site of action potential initiation and is necessary for reproducing the sharp spike-onset kink characteristic of fast-spiking interneurons [24]. In this two-compartment HH model, the soma and AIS are connected by an axial conductance *g*_*c*_, permitting current flow between compartments. All injected currents and the stochastic OU noise process were applied to the soma, matching the standard whole-cell recording configuration in which intrinsic noise is measured at the soma [25].

**Fig. 2.**
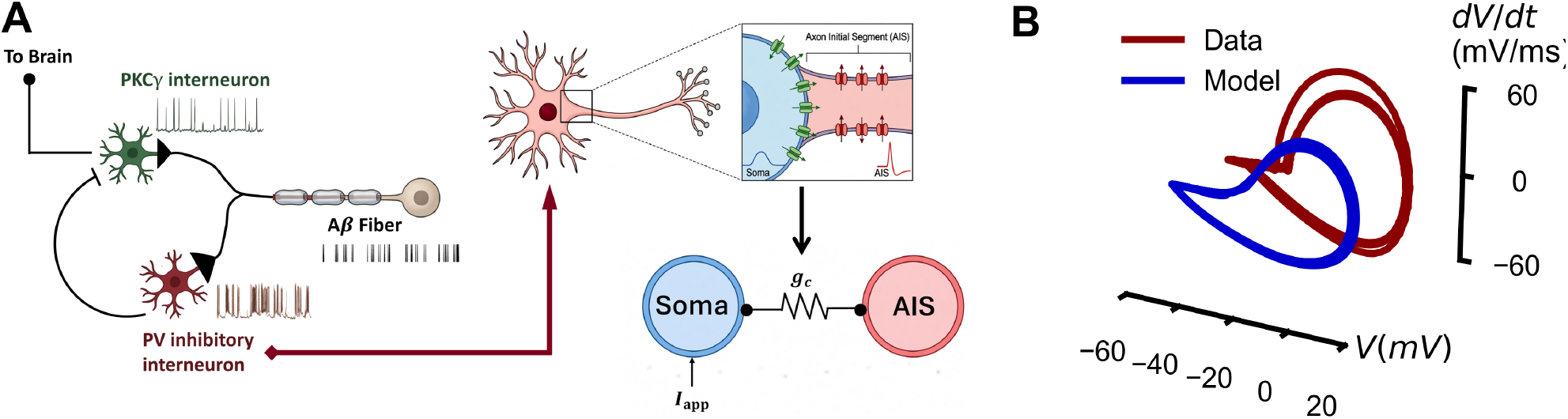
Two-compartment Hodgkin–Huxley (HH) model and its validation against experimental data. (A) Schematic of the dorsal horn neural circuit (left), illustrating the established connectivity within laminae II–III, where A*β* afferents synapse onto both PVINs and PKC*γ* interneurons, while PVINs provide feed-forward inhibition to PKC*γ* interneurons, which project to higher-order brain regions. spPVINs, like quiescent PVINs, serve as inhibitory interneurons within this circuit. Anatomy of spPVINs (right) with a zoomed-in inset, illustrating the two compartments: the axon initial segment (AIS) and soma. Two-compartment HH model, showing the coupling between the soma and AIS with conductance *g*_*c*_ (bottom right). (B) Phase-plane plot of action potential cycles generated by a phasic spPVIN (the same neuron as the one highlighted in Figure 1A, fourth panel) alongside the fitted action potential cycle generated by the two-compartment HH model (blue).

The somatic membrane voltage *V* is governed by the equation

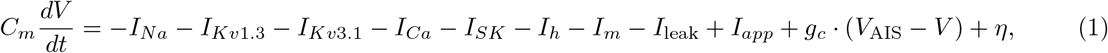

where *C*_*m*_ is the membrane capacitance, *I*_*j*_ are the somatic ionic currents (*j* = *Na, Kv*1.3, *Kv*3.1, *Ca, SK, h, m*, leak), *I*_*app*_ is the applied current, *η* is the OU noise process, and *g*_*c*_*·*(*V*_AIS_*−V*) is the axial coupling current from the AIS with coupling conductance *g*_*c*_. The AIS voltage equation (*V*_AIS_), on the other hand, satisfies

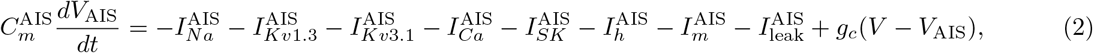

where 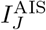 are the ionic currents in the AIS (*J* = *Na, Kv*1.3, *Kv*3.1, *Ca, SK, h, m*, leak), and 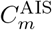 is the AIS membrane capacitance, obtained by scaling the somatic capacitance *C*_*m*_ by the compartment area ratio *κ* (Table 1). Note that SK channels were not included in the AIS (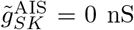; Table 1), so that 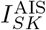 vanishes under all conditions considered here. The choice of currents was based on [8] and extended to additionally incorporate *I*_*h*_ (HCN channel current) and *I*_*m*_ (Kv7 or M-current).

**Table 1.** Model Parameters for the Two-Compartment Hodgkin–Huxley spPVIN Model.

| Parameter | Value | Description |
| --- | --- | --- |
| **Global Parameters & Reversal Potentials** |  |  |
| $C_m$ | 30 pF | Membrane capacitance |
| $E_{Na}$ | 49 mV | Na<sup>+</sup> Nernst potential |
| $E_K$ | -88 mV | K<sup>+</sup> Nernst potential |
| $E_{Ca}$ | 63 mV | Ca<sup>2+</sup> Nernst potential |
| $E_{leak}$ | -60 mV | Leak reversal potential |
| $E_h$ | -22 mV | $h$-current reversal potential |
| **Somatic Conductances** |  |  |
| $\tilde{g}_{Na}$ | 106.2 nS | Maximum transient Na<sup>+</sup> conductance |
| $\tilde{g}_{Kv1}$ | 18 nS | Maximum delayed-rectifier K<sup>+</sup> (Kv1.3) conductance |
| $\tilde{g}_{Kv3}$ | 72 nS | Maximum fast delayed-rectifier K<sup>+</sup> (Kv3.1) conductance |
| $\tilde{g}_h$ | 9 nS | Maximum hyperpolarization-activated ($h$-) conductance |
| $f_0$ | 0.6 | Proportion of fast component in somatic $h$-current |
| $\tilde{g}_m$ | 4.5 nS | Maximum M-current conductance |
| $\tilde{g}_{SK}$ | 9 nS | Maximum SK current conductance |
| $\tilde{g}_{Ca}$ | 5.4 nS | Maximum high-voltage activated Ca<sup>2+</sup> conductance |
| $\tilde{g}_{leak}$ | 3.6 nS | Maximum leak conductance |
| **Axon Initial Segment (AIS) Conductances** |  |  |
| $\tilde{g}_{Na}^{AIS}$ | 11.8 nS | Maximum transient Na<sup>+</sup> conductance (AIS) |
| $\tilde{g}_{Kv1}^{AIS}$ | 2 nS | Maximum delayed-rectifier K<sup>+</sup> (Kv1.3) conductance (AIS) |
| $\tilde{g}_{Kv3}^{AIS}$ | 8 nS | Maximum fast delayed-rectifier K<sup>+</sup> (Kv3.1) conductance (AIS) |
| $\tilde{g}_h^{AIS}$ | 1 nS | Maximum hyperpolarization-activated ($h$-) conductance (AIS) |
| $f_0^{AIS}$ | 0.6 | Proportion of fast component in AIS $h$-current |
| $\tilde{g}_m^{AIS}$ | 0.5 nS | Maximum M-current conductance (AIS) |
| $\tilde{g}_{SK}^{AIS}$ | 0 nS | Maximum SK current conductance (AIS) |
| $\tilde{g}_{Ca}^{AIS}$ | 0.6 nS | Maximum high-voltage activated Ca<sup>2+</sup> conductance (AIS) |
| $\tilde{g}_{leak}^{AIS}$ | 0.4 nS | Maximum leak conductance (AIS) |
| **Ca<sup>2+</sup> Dynamics & Coupling** |  |  |
| $A$ | 2700 $\mu\text{m}^2$ | Cell surface area |
| $d$ | 0.1 $\mu\text{m}$ | Shell thickness for Ca<sup>2+</sup> dynamics |
| $[\text{Ca}^{2+}]_r$ | 0.07 $\mu\text{M}$ | Resting intracellular Ca<sup>2+</sup> concentration |
| $B_{tot}$ | 92 $\mu\text{M}$ | Total Ca<sup>2+</sup> buffer concentration |
| $K_d$ | 0.1 $\mu\text{M}$ | Buffer affinity/dissociation constant |
| $\gamma$ | 0.01 $\text{ms}^{-1}$ | Ca<sup>2+</sup> recovery rate |
| $k_{SK}$ | 0.8 $\mu\text{M}$ | Ca<sup>2+</sup> sensitivity (half-activation) of SK channels |
| $F$ | 0.096485 C $\mu\text{mol}^{-1}$ | Faraday constant scaling factor |
| $g_c$ | 10 nS | Coupling conductance between soma and AIS |
| $\kappa$ | 0.9 | Compartment area ratio (soma to total) |
| $P_{NMDA}^{\text{Ca}^{2+}}$ | 0.1 | Permeability fraction of Ca<sup>2+</sup> influx due to the NMDA current |
| $P_{AMPA}^{\text{Ca}^{2+}}$ | 0.03 | Permeability fraction of Ca<sup>2+</sup> influx due to the AMPA current |

Each ionic current included in the spPVIN model takes the canonical conductance-based formalism, given by 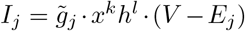, where 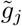 is the maximal conductance, *x, h* are the (in)activation gating variables, *k, l* are the number of gates of the (in)activation variables, and *E*_*j*_ is the reversal potential of a particular current.

Seven distinct ionic currents were incorporated into the soma (Eq. (1)) and AIS (Eq. (2)). These include:

1. **Hyperpolarization-activated current (***I*_*h*_), given by

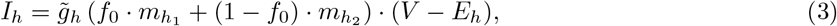

where 0 *≤ f*_0_ *≤* 1 and 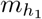 and 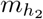 are, respectively, the fast and slow activation variables of *I*_*h*_ whose dynamics are governed by

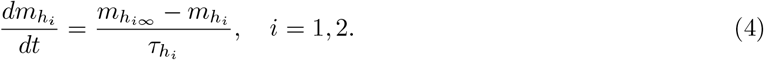

The time constant and steady state activation of 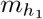 are given by

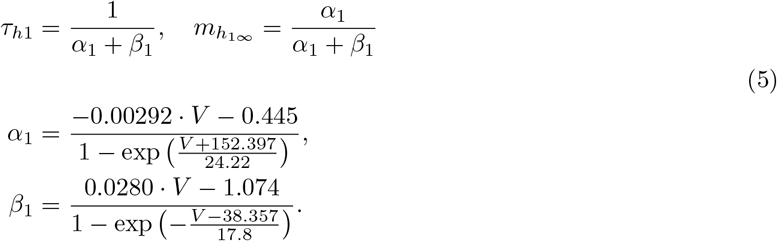

Similarly, the time constant and steady state activation of 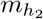 are given by

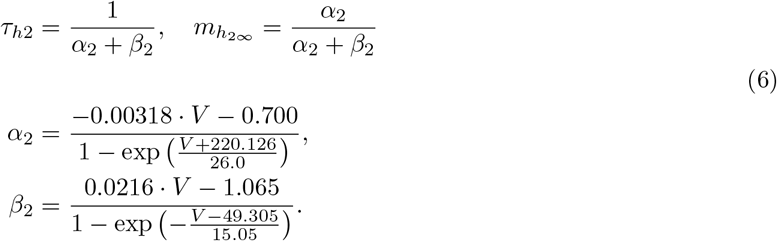
2. **Fast transient Na**^+^ **current (***I*_*Na*_), described by

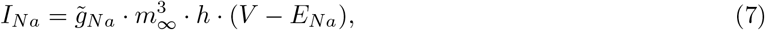

where *m*_*∞*_ is the steady state activation, given by

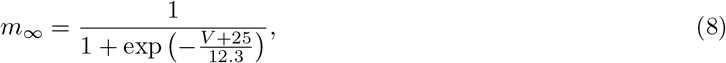

and *h* is the inactivation variable, satisfying

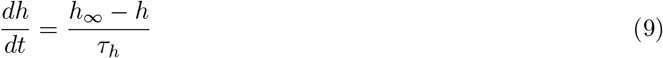

with time constant *τ*_*h*_ and steady state inactivation *h*_*∞*_, given by

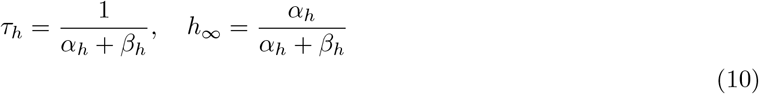

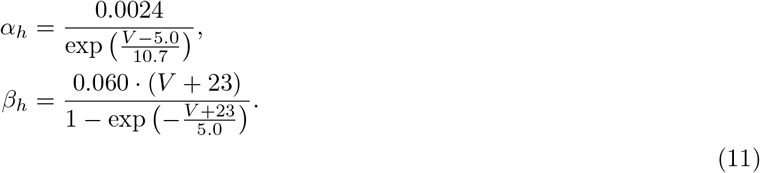
3. **Slow Kv1.3-mediated delayed-rectifier K**^+^ **current (***I*_*Kv*1.3_), defined by

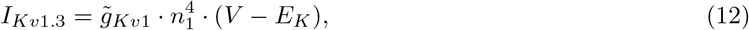

where *n*_1_ is the activation variable, satisfying

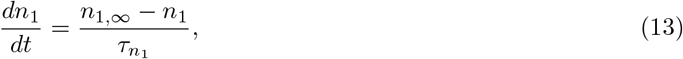

with time constant 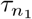 and steady state activation *n*_1,*∞*_, given by

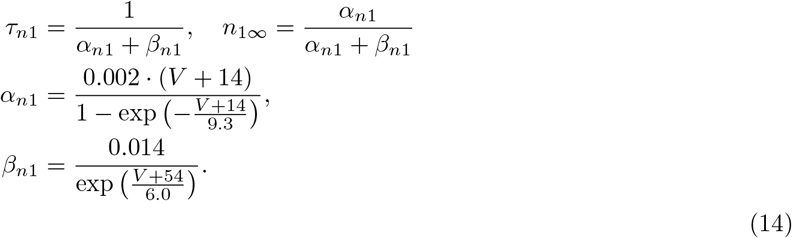
4. **Fast Kv3.1-mediated delayed-rectifier K**^+^ **current (***I*_*Kv*3.1_), expressed as

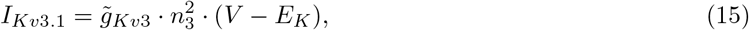

where *n*_3_ is the activation variable, satisfying

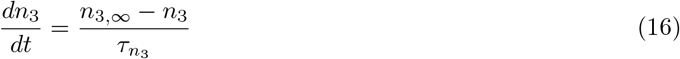

with time constant 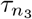 and steady state activation *n*_3,*∞*_, given by

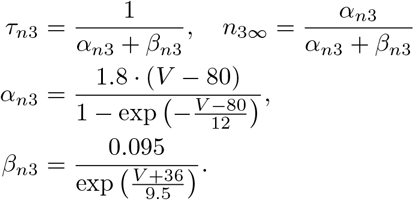
5. **Slow, non-inactivating M-type K**^+^ **current (***I*_*m*_), described by

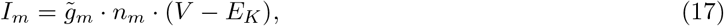

where *n*_*m*_ is the activation variable, satisfying

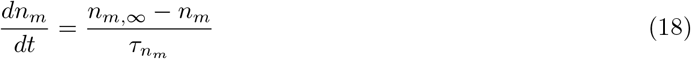

with time constant 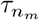 and steady state activation *n*_*m,∞*_, given by

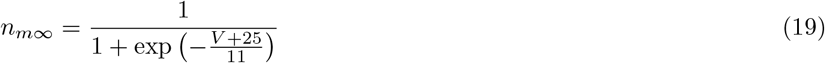

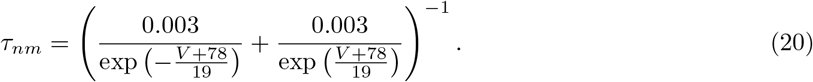
6. **High-voltage-activated Ca**^2+^ **current (***I*_*Ca*_), expressed as

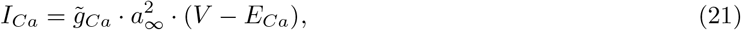

where *a*_*∞*_ is the steady state activation, given by

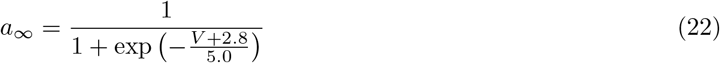
7. **Ca**^2+^**-dependent K**^+^ **current (***I*_*SK*_), defined by

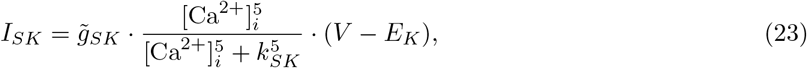

where [Ca^2+^]_*i*_ denotes intracellular Ca^2+^ concentration. Its dynamics are governed by both transmembrane influx and clearance through buffering and extrusion, as described by the equation

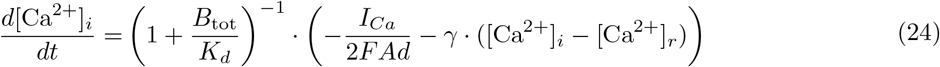

where *B*_tot_ is the total buffer concentration, *K*_*d*_ is the dissociation constant of the buffer, *γ* is the extrusion rate, and [Ca^2+^]_*r*_ is the resting Ca^2+^ concentration.

An exponentially correlated noise process in the form of OU noise (*η*) is incorporated into the model to capture the stochastic fluctuations in the membrane voltage. Its dynamics are governed by the equation

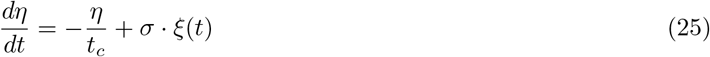

where *t*_*c*_ sets the correlation timescale, 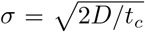 *D* is the noise intensity), and *ξ*(*t*) is a zero-mean unit-variance white Gaussian noise process [26, 27]. With *V* expressed in mV and *t*_*c*_ in ms, *σ* has units of mV ms^*−*1*/*2^.

#### Electrophysiological feature extraction

To quantify the electrophysiological properties of the two-compartment model, six distinct metrics were extracted from the simulated somatic membrane potential (*V*) and intracellular calcium concentration ([Ca^2+^]_*i*_) traces. For spike-dependent metrics, action potentials were detected whenever the somatic membrane potential crossed *−* 20 mV from below (the same spike-detection criterion used in the circuit simulations) and were isolated using a 6 ms window (*±*3 ms) centered on each detected crossing. All features were also averaged over *N* = 10 realizations, and each realization was simulated for 90 s. The six features quantified include:

1. *Mean firing rate*. Calculated as the total number of detected action potentials divided by the total duration of the simulation, expressed in Hz.
2. *Resting membrane potential*. Computed as the median of the membrane potential during a quiescent period (guaranteeing a robust measure that is insensitive to transient spikes and to the alternation between bursting and quiescent states).
3. *Mean intracellular calcium*. Computed as the time-averaged mean of the [Ca^2+^]_*i*_ trace over the entire simulation period.
4. *Spike amplitude*. Calculated as the voltage difference between the peak of each detected action potential and baseline (defined as the local minimum of the membrane potential immediately preceding the spike peak within the local analysis window).
5. *Spike half-width*. Computed as the temporal duration (in ms) during which the action potential remains above its half-maximal voltage (defined as the arithmetic midpoint between baseline voltage and spike peak).
6. *Spike threshold*. Quantified as the somatic membrane potential (*V*) at the exact time point during the action potential upstroke when the rate of membrane depolarization (*dV/dt*) first reaches or exceeds 10 mV/ms.

For all spike features, including amplitude, half-width and threshold, the final reported values for each realization represent the arithmetic mean of the respective metrics averaged across all valid individual spikes detected during the simulation.

#### Circuit design

To investigate the circuit-level properties of spPVINs, we constructed a dorsal horn circuit model (Figure 2A) based on the architecture described previously [8, 28]. The model captures the established connectivity within laminae II–III of the spinal dorsal horn, in which A*β* afferents synapse onto both PVINs and PKC*γ* interneurons, while PVINs provide GABAergic/glycinergic feed-forward inhibition onto PKC*γ* cells [29–31]. Parameters of the PKC*γ* HH model are listed in Table 2, and its equations are given in the Supplementary Material.

**Table 2.** PKC*γ* Model Parameters [28].

| Parameter | Value | Unit | Definition |
| --- | --- | --- | --- |
| $C_{m,\gamma}$ | 1 | $\mu\text{F}/\text{cm}^2$ | Specific membrane capacitance |
| $\tilde{g}_{Na1}$ | 0.1625 | $\text{mS}/\text{cm}^2$ | Maximum HH-type 1 $\text{Na}^+$ conductance |
| $\tilde{g}_{Na2}$ | 85.48 | $\text{mS}/\text{cm}^2$ | Maximum HH-type 2 $\text{Na}^+$ conductance |
| $\tilde{g}_K$ | 4.3 | $\text{mS}/\text{cm}^2$ | Maximum HH-type $\text{K}^+$ conductance |
| $\tilde{g}_A$ | 10.9 | $\text{mS}/\text{cm}^2$ | Maximum A-type $\text{K}^+$ conductance |
| $\tilde{g}_{KDR}$ | 3.111 | $\text{mS}/\text{cm}^2$ | Maximum delayed-rectifier $\text{K}^+$ conductance |
| $\tilde{g}_{leak}$ | 0.96 | $\text{mS}/\text{cm}^2$ | Maximum leak conductance |
| $E_{Na}$ | 50 | mV | $\text{Na}^+$ Nernst potential |
| $E_K$ | -70 | mV | $\text{K}^+$ Nernst potential |
| $E_{leak}$ | -65 | mV | Leak Nernst potential |
| $L$ | 20 | $\mu\text{m}$ | Soma length |
| $D_{soma}$ | 20 | $\mu\text{m}$ | Soma diameter |
| $T$ | 23 | $^{\circ}\text{C}$ | Temperature |

Excitatory A*β* afferent input was modeled as a Poisson process generating independent spike trains at specified constant stimulation frequencies (*f*_*Aβ*_ = 0–15 Hz). Each simulated A*β* spike simultaneously triggered AMPA and NMDA conductance transients in both the spPVIN HH model and the PKC*γ* HH model (described in the Supplementary Material). For circuit simulations, the Ornstein–Uhlenbeck (OU) noise *η* was removed from the spPVIN HH model and replaced by a synaptic current *I*_*syn*_ representing the modeled synaptic inputs. To close the feed-forward inhibitory loop, spPVIN action potentials were detected dynamically during integration whenever the PVIN somatic membrane potential crossed a threshold of *−*20 mV from below. Each detected PVIN spike subsequently triggered glycine and GABA_*A*_ conductance transients in the downstream PKC*γ* HH model.

Excitatory and inhibitory synaptic conductances incorporated into both the spPVIN and PKC*γ* HH models were modeled using dual-exponential kinetics, with conductance

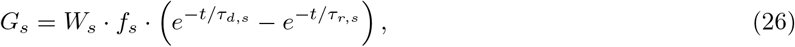

where *τ*_*r,s*_ and *τ*_*d,s*_ are the rise and decay time constants, *W*_*s*_ is the absolute lumped synaptic weight (in nS), and *f*_*s*_ normalizes the peak conductance such that the maximum value of *G*_*s*_(*t*) equals *W*_*s*_ [32]. Excitatory A*β* inputs into the spPVIN and PKC*γ* HH models were mediated by AMPA and NMDA receptors (*E*_*exc*_ = 0 mV). The NMDA conductance included a voltage-dependent Mg^2+^ block [33], described by

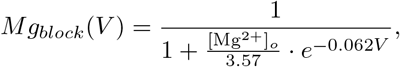

with [Mg^2+^]_*o*_ = 1 mM. Inhibitory spPVIN*→*PKC*γ* synapses used co-released glycine and GABA_*A*_ receptors with *E*_*inh*_ = *−* 70 mV [29]. Synaptic weights and kinetic parameters are detailed in Table 3.

**Table 3.** Circuit Synaptic Weights and Kinetics.

| Connection | Receptor | Weight ($W_s$) | $\tau_r$ (ms) | $\tau_d$ (ms) |
| --- | --- | --- | --- | --- |
| $A\beta \rightarrow \text{PVIN}$ | AMPA | 2.0 nS | 0.1 | 5.0 |
| $A\beta \rightarrow \text{PVIN}$ | NMDA | 3.0 nS | 2.0 | 100.0 |
| $A\beta \rightarrow \text{PKC}\gamma$ | AMPA | 5.5 nS | 0.1 | 5.0 |
| $A\beta \rightarrow \text{PKC}\gamma$ | NMDA | 2.0 nS | 2.0 | 100.0 |
| $\text{PVIN} \rightarrow \text{PKC}\gamma$ | Glycine | 5.32 nS | 0.1 | 10.0 |
| $\text{PVIN} \rightarrow \text{PKC}\gamma$ | GABA<sub>A</sub> | 10.0 nS | 0.1 | 20.0 |

Given that AMPA and NMDA currents contribute to the synaptic current and permit Ca^2+^ influx, the Ca^2+^ flux-balance equation for the spPVIN HH model must account for this additional source of Ca^2+^ influx. Accordingly, the Ca^2+^ balance equation becomes

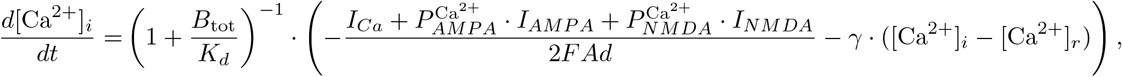

where 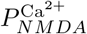 and 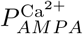 are the permeability fractions of Ca^2+^ influx due to the NMDA and AMPA currents, respectively.

To quantify the effects of spPVIN feed-forward inhibition on PKC*γ* responses within the circuit, we used two complementary measures:

1. *Circuit gain* (*CG*), defined as the linear-regression slope of the trial-averaged PKC*γ* firing rate against the A*β* input frequency over the 0–15 Hz range. To avoid underestimation from the flat sub-threshold region, the fit was restricted to points where the PKC*γ* rate exceeded 0.5 Hz; if fewer than three points met this criterion, the entire frequency-response curve was used. Gain is expressed in Hz PKC*γ* per Hz A*β*.
2. *Suppression ratio* (*SR*), defined as

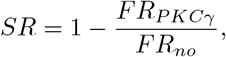

where *FR*_*no*_ denotes the firing rate of PKC*γ* neurons in the absence of feed-forward inhibition.

Both measures were computed for each simulation and averaged across *N* = 10 independent realizations of the circuit. Each realization lasted 2.5 s, comprising a 500-ms silent period allowing the circuit to reach steady state, followed by a 2-s epoch of A*β* synaptic input.

#### Model fitting and numerical implementation

The two-compartment HH model of spPVINs was integrated in Python. The model was parameterized by fitting the simulated action potential cycle generated by the model to the action potential cycle of a phasic-firing spPVIN (Figure 2B). Fitting the action potential cycle ensures that the model accurately captures both the action potential waveform and the kinetics of the underlying ion channels.

To characterize the intrinsic (deterministic) dynamics of the resulting HH model, we removed noise and computed the bifurcation diagrams with respect to different parameters of the model, including *I*_app_, 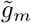 and 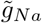. Using the same model, we conducted a slow-fast analysis by treating the Ca^2+^ variable as a parameter and analyzing the resulting bifurcation structures of the model with respect to this new parameter. All bifurcation analyses were performed using XPPAUT and AUTO-07p, both freely available software packages (XPPAUT: https://sites.pitt.edu/~phase/bard/bardware/xpp/xpp.html; AUTO-07p: https://github.com/auto-07p/auto-07p).

For the circuit simulations, we performed 10 independent trials for each condition to capture stochastic variability in spike timing. A*β* input was delivered as Poisson spike trains with mean firing rates ranging from 0 to 15 Hz in 1-Hz increments. Each frequency epoch lasted 2 s and was preceded by a 500-ms silent period to allow the circuit to reach a steady state before stimulation.

All Python code and XPPAUT files used to analyze the model and the circuit numerically are available in a GitHub repository (https://github.com/rfritzdj/spPVIN). Model parameters are listed in Table 1.

## Results

### Underlying dynamics of the two-compartment HH model

To examine the mechanisms underlying the diverse firing phenotypes of spPVINs, we first characterized the model’s excitability and dynamical regimes as a function of the applied current (*I*_*app*_).

To this end, we plotted the bifurcation diagram of the somatic membrane voltage (*V*) with respect to *I*_*app*_ (Figure 3A). The diagram revealed the presence of three distinct dynamic regimes, two of which are quiescent at low and high *I*_*app*_ and one that is oscillatory for intermediate *I*_*app*_ (Figure 3A). At low *I*_*app*_, a branch of stable equilibria exhibiting type III excitability is obtained. As *I*_*app*_ increases, this branch of equilibria loses stability by undergoing a subcritical Hopf bifurcation (HB1), forming a branch of unstable equilibria. Meanwhile, two envelopes of unstable limit cycles emerge from HB1. Increasing *I*_*app*_ further causes the branch of unstable equilibria to regain stability through a second subcritical Hopf bifurcation (HB2), forming a branch of stable equilibria representing the depolarization block. As was the case for HB1, two envelopes of unstable limit cycles also emerge from HB2. These envelopes eventually undergo saddle-node bifurcations of periodic orbits (SNP1 at low *I*_*app*_ and SNP2 at high *I*_*app*_, respectively), with the former giving rise to an envelope of stable bursting orbits and the latter to an envelope of stable limit cycles. These two envelopes meet at a torus bifurcation point (TR) at intermediate *I*_*app*_, reflecting the birth of mixed-mode oscillations at lower *I*_*app*_. The subcritical nature of HB1 and HB2 gives rise to two bistable regimes, between SNP1 and HB1 and between HB2 and SNP2, consistent with type II excitability. Taken together, these results provide an overview of the distinct patterns of dynamical behavior that spPVINs can exhibit according to the HH model. In particular, they demonstrate that spPVINs can display both bursting (Figure 3B, top) and tonic firing (Figure 3B, bottom) activity, revealing key information about their electrical properties.

**Fig. 3.**
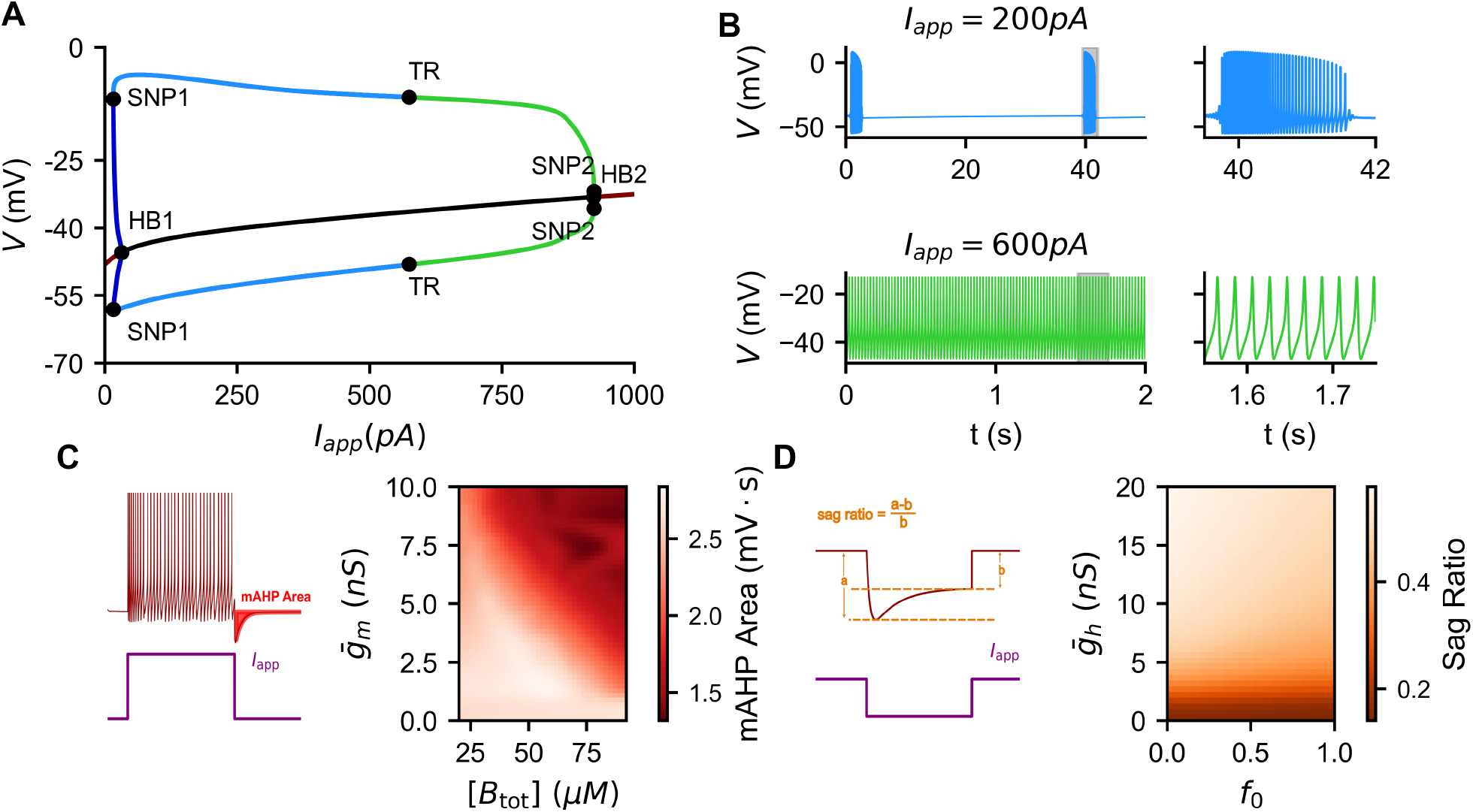
Two-compartment Hodgkin–Huxley (HH) model reproduces key features of neuronal excitability of spPVINs. (A) Bifurcation diagram of somatic membrane potential *V* as a function of applied current *I*_*app*_; red (black) lines: branch of stable (unstable) equilibria; green (blue) lines: envelopes of stable (unstable) limit cycles; cyan lines: envelopes of stable bursting orbits. HB1, HB2: Hopf bifurcations; SNP1, SNP2: saddle-node bifurcations of periodic orbits; TR: torus bifurcation. (B) Sample voltage traces at *I*_*app*_ = 200 pA and *I*_*app*_ = 600 pA showing bursting and tonic firing activity, respectively. (C) Medium afterhyperpolarization (mAHP) following the termination of an injected current pulse (left) and heatmap illustrating the effects of the maximum conductance of the M-type K^+^ current 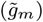 and the total intracellular buffer concentration (*B*_tot_) on mAHP area following injection of a depolarizing current of *I*_*app*_ = 150 pA (right). The heatmap is color-coded according to the scale shown in the color bar on the right. (D) Sag response evoked by injection of a hyperpolarizing current, mediated by the hyperpolarization-activated current *I*_*h*_ (left); the sag response is quantified using the sag ratio. Heatmap illustrating the effects of the maximum conductance of 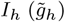 and the fractional conductance contribution of the fast *I*_*h*_ activation component (*f*_0_) on the sag ratio (right). The heatmap is color-coded according to the scale shown in the color bar on the right.

The HH model includes two subthreshold currents that together shape the excitability of spPVINs: a slowly activating, non-inactivating M-type K^+^ current (*I*_*m*_) that opposes membrane depolarization and underlies spike-frequency adaptation [34, 35], and the hyperpolarization-activated cation (HCN) current (*I*_*h*_) recruited during membrane hyperpolarization, which produces an inward depolarizing drive that generates voltage sag and rebound responses [36, 37].

To explore the contribution of each of these two currents to the deterministic dynamics of the HH model (in the absence of noise), we first investigated the effects of *I*_*m*_ on the medium afterhyperpolarization (mAHP). The mAHP was defined as the hyperpolarizing voltage deflection occurring between the end of the current step and 200 ms thereafter [35, 38, 39]. The mAHP area (Figure 3C, left) was quantified as the area between the resting membrane potential and the mAHP voltage trajectory over the same 200-ms interval. A larger mAHP area typically produces stronger spike-frequency adaptation and lengthens interspike intervals, whereas a smaller mAHP area permits higher firing rates [38, 40]. In the HH model of spPVINs, increasing the maximum conductance of 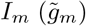 reduces mAHP area when a depolarizing current is injected, an effect that is diminished by increasing the total intracellular buffer concentration *B*_tot_ (Figure 3C, right). This action of 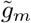 on the mAHP is indirect: because the mAHP in this model is generated largely by *I*_*SK*_, a larger 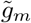 limits firing during the depolarizing step, reduces cumulative Ca^2+^ influx, and therefore weakens *I*_*SK*_ activation. Consistently, increasing *B*_tot_ attenuates intracellular Ca^2+^ transients, thereby also reducing the activation of *I*_*SK*_ and weakening the hyperpolarizing drive that shapes the mAHP.

We next examined *I*_*h*_, which contributes primarily to subthreshold excitability [36, 37] and is recruited in response to hyperpolarization. We quantified its contribution using the sag ratio, defined as the normalized voltage sag during a hyperpolarizing current step,

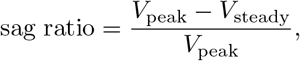

where *V*_peak_ and *V*_steady_ are the peak and steady-state voltage deflections relative to rest, respectively (Figure 3D, left). A larger sag ratio reflects stronger *I*_*h*_ activation, which partially offsets the hyperpolarization and accelerates the return to resting potential. As expected, when a hyperpolarizing current of *I*_*app*_ = *−* 150 pA was injected, the HH model of spPVINs showed a monotonic increase in sag ratio as the maximum conductance of 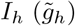 was increased (Figure 3D, right), confirming that *I*_*h*_ is the primary determinant of voltage sag. The relative weighting of fast and slow *I*_*h*_ activation components (*f*_0_) has little to no effect on the sag ratio, indicating that it is the total *I*_*h*_ conductance rather than its kinetics that governs this feature.

### Bifurcation analysis reveals an intrinsic bursting mechanism defined by *I*_***m***_ **and *I***_***Na***_

To determine whether the two-compartment spPVIN HH model possesses an intrinsic mechanism for bursting, we performed numerical continuation in the absence of applied current (*I*_app_ = 0 pA), focusing on the maximum conductances of the 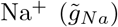, M-type 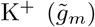, and HCN 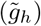 currents. We asked whether varying these conductances alone could drive the model from a quiescent state into a bursting regime, thereby revealing how these intrinsic currents interact to govern spontaneous activity.

We first examined the effects of 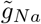 and 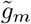 independently by plotting bifurcation diagrams of *V* with respect to each conductance at *I*_app_ = 0 pA (Figure 4A and B, respectively). Varying 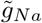 revealed a transition from quiescence to intrinsic bursting (Figure 4A). Indeed, at low 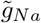, the model produces a branch of stable equilibria (with type III excitability, which permits the generation of isolated spikes). As 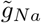 increases, this branch loses stability through a subcritical Hopf bifurcation (HB_*Na*_), giving rise to envelopes of unstable limit cycles. Further increases in 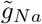 lead these envelopes to undergo a saddle-node bifurcation of periodic orbits (SNP_*Na*_), beyond which envelopes of stable bursting orbits emerge. These bursting envelopes subsequently undergo two period-doubling bifurcations (PD_*Na*,1_ and PD_*Na*,2_), indicating the emergence of bursting orbits (similar to the representative bursting trajectory seen in the inset, Figure 4A) with an increasing number of spikes per burst. Interestingly, within this structure, a regime of bistability, or coexistence between quiescent and bursting states, is formed between SNP_*Na*_ and HB_*Na*_.

**Fig. 4.**
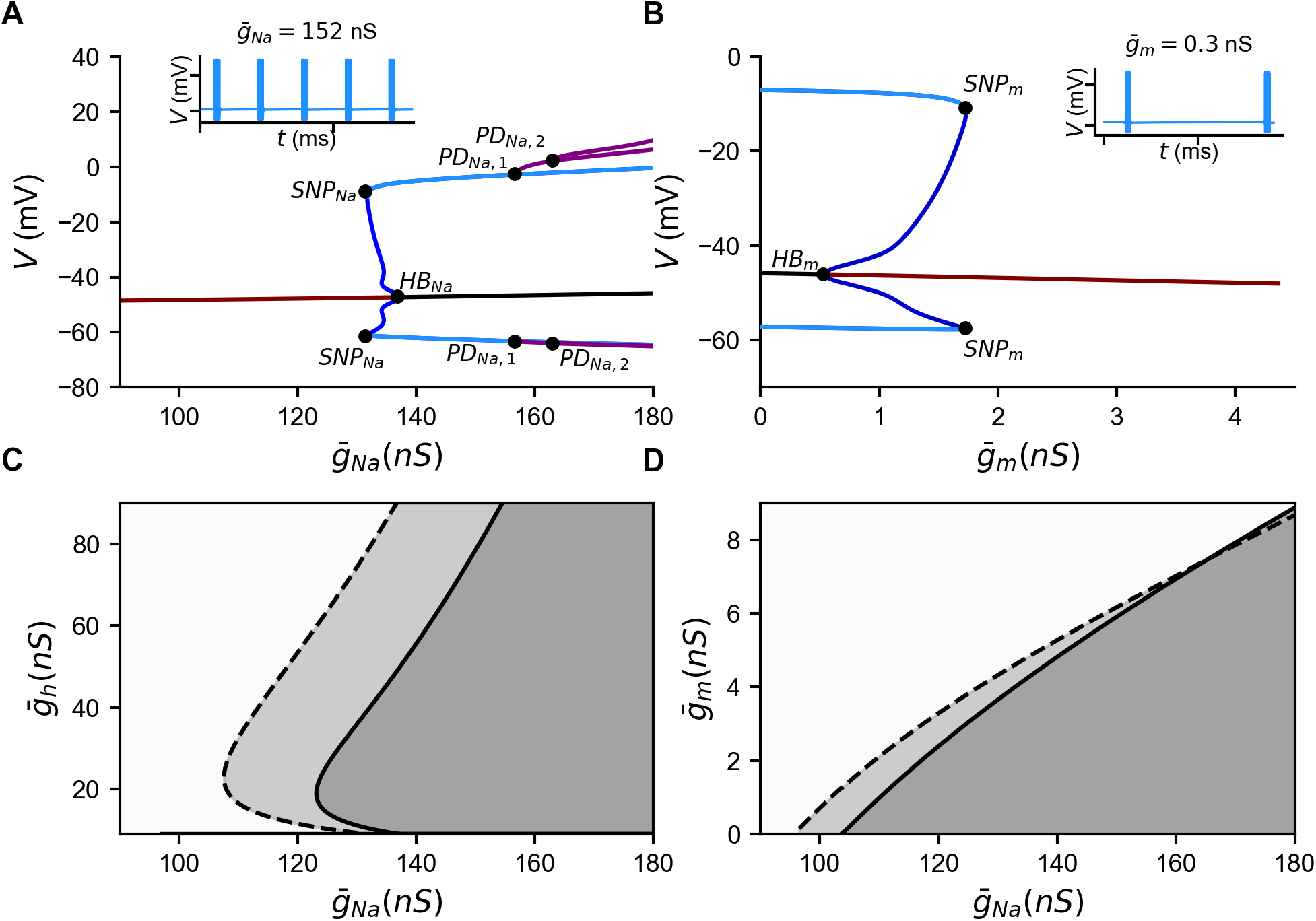
Intrinsic bursting in the Hodgkin–Huxley (HH) model of spPVINs is shaped by the maximum conductances of the 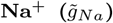, M-type 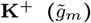, and hyperpolarization-activated 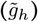 currents. (A,B) Bifurcation diagrams of somatic membrane potential *V* with respect to 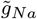 (A), and 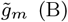 at zero applied current (*I*_app_ = 0 pA). Red (black) lines: branch of stable (unstable) equilibria; blue lines: envelopes of unstable limit cycles; cyan lines: envelopes of stable bursting orbits; purple lines: period-doubling branches. HB_*Na*_, HB_*m*_: Hopf bifurcations; SNP_*Na*_, SNP_*m*_: saddle-node bifurcations of periodic orbits; PD_*Na*,1_, PD_*Na*,2_: period-doubling bifurcations. Insets: representative time courses of *V* for bursting orbits at 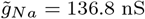 in A, and 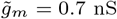 in B. (C,D) Two-parameter bifurcation diagrams of the HH model with respect to 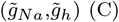, and 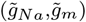. Solid lines correspond to the Hopf bifurcations: HB_*Na*_ (C) and HB_*m*_ (D), while dashed lines correspond to the saddle-node bifurcations of periodic orbits: SNP_*Na*_ (C) and SNP_*m*_ (D). White: quiescent regime; light gray: bistability regime; dark gray: bursting regime.

In contrast, varying 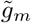 produces the opposite effect (Figure 4B, with a representative bursting trajectory shown in the inset). At low 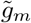, the model exhibits envelopes of stable bursting orbits surrounding a branch of unstable equilibria. As 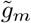 increases, these envelopes terminate at a saddle-node bifurcation of periodic orbits (SNP_*m*_), from which two envelopes of unstable periodic orbits emerge and subsequently terminate at a subcritical Hopf bifurcation (HB_*m*_). The branch of unstable equilibria gains stability at HB_*m*_, generating a branch of stable equilibria. Once again, a bistable regime emerges between SNP_*m*_ and HB_*m*_, beyond which bursting is eliminated, leaving the quiescent state as the sole stable state at high 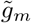 values. These results thus show that 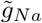 promotes intrinsic bursting, whereas increasing 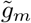 opposes it, indicating that the balance between these fast inward and slow outward currents is a key determinant of intrinsic bursting [16].

We next examined how 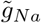 interacts with 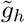 and 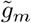 using two-parameter bifurcation analysis. Specifically, we constructed bifurcation diagrams in the 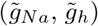 ane (Figure 4C) and in the 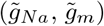 plane (Figure 4D) by tracking both Hopf bifurcations and saddle-node bifurcations of periodic orbits. In the 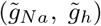-plane, increasing 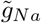 promotes a transition from quiescence (where isolated spikes can be triggered) through bistability into bursting, whereas 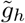 shifts the bifurcation boundaries toward higher 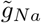 in a biphasic manner, particularly at low and high 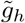, thereby narrowing the parameter range that supports bursting (Figure 4C). In the 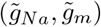 plane, on the other hand, increasing 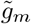 shifts the bifurcation boundaries monotonically toward higher 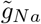, reflecting its antagonistic interaction with 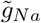 (Figure 4D). The bistable regime also progressively shrinks and eventually disappears at high 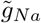 and 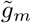, where the subcritical Hopf bifurcation becomes supercritical. Taken together, these results reveal a mechanism for both isolated spiking and intrinsic bursting, governed by the interplay among these three maximal conductances. Together, they shape a dynamical landscape in which stochastic fluctuations arising from channel and/or synaptic noise can drive the system between quiescence, isolated spiking, and bursting. This provides a potential mechanism by which the HH model can intrinsically generate stochastic transitions between these activity states, consistent with the experimentally observed irregular firing patterns (Figure 1A).

### *I*_*h*_-mediated noise-driven spontaneous firing in spPVINs

As we have seen above (Figure 1A), most PVINs are quiescent in the absence of injected current, but a small subset (spPVINs) exhibits spontaneous irregular firing, in the form of isolated spikes and bursts. Furthermore, our bifurcation analysis demonstrated that the spPVIN HH model can intrinsically generate bursting activity (Figure 4). We therefore sought to investigate how stochastic fluctuations in membrane potential may contribute to the transitions between quiescence, isolated spiking and spontaneous burst firing. In particular, we examined whether noise can stochastically drive the model between these three states, and how this process depends on *I*_*h*_.

To do so, we simulated the HH model under two different conditions, in the absence and presence of *I*_*h*_, while systematically varying the noise parameters, including noise amplitude (*σ*) and temporal correlation timescale (*t*_*c*_). Three distinct values of *t*_*c*_ (1, 10 and 100 ms) and a whole range of *σ* (between 0.01–0.1 mV ms^*−*1*/*2^; adjusted based on the noise intensity *D*) were tested. Our results demonstrated that in the absence of *I*_*h*_, both *σ* and *t*_*c*_ need to be relatively large to reliably elicit action potentials (Figure 5A, B). Indeed, the vast majority of simulated traces remain silent, resembling the activity of quiescent PVINs. Isolated action potentials are observed only at the longest *t*_*c*_ tested and at a relatively high *σ* (*σ* = 0.07 mV ms^*−*1*/*2^), whereas further increasing *σ* induces bursting. This suggests that quiescent PVINs may express low levels of functional HCN channels, resulting in a weak *I*_*h*_-mediated depolarizing drive, and/or may be less susceptible to synaptic fluctuations. In contrast, the presence of *I*_*h*_ substantially lowers the noise requirements for spontaneous firing (Figure 5C, D). Isolated action potentials emerge at lower *t*_*c*_ values, with firing observed at *t*_*c*_ = 10 ms for *σ* as low as 0.05 mV ms^*−*1*/*2^, while at *t*_*c*_ = 100 ms, isolated spikes can be elicited with *σ* as low as 0.02 mV ms^*−*1*/*2^. Increasing either *σ* or *t*_*c*_ progressively shifts the firing pattern from isolated spiking toward bursting. Interestingly, increasing both *σ* and *t*_*c*_ increases the firing rate (Figure 5D), indicating that larger and more persistent fluctuations provide a stronger and more sustained depolarizing drive. The longer persistence of these fluctuations allows the membrane potential to remain near threshold for longer periods, facilitating the recruitment of slow intrinsic currents and promoting the transition from isolated spiking to bursting.

**Fig. 5.**
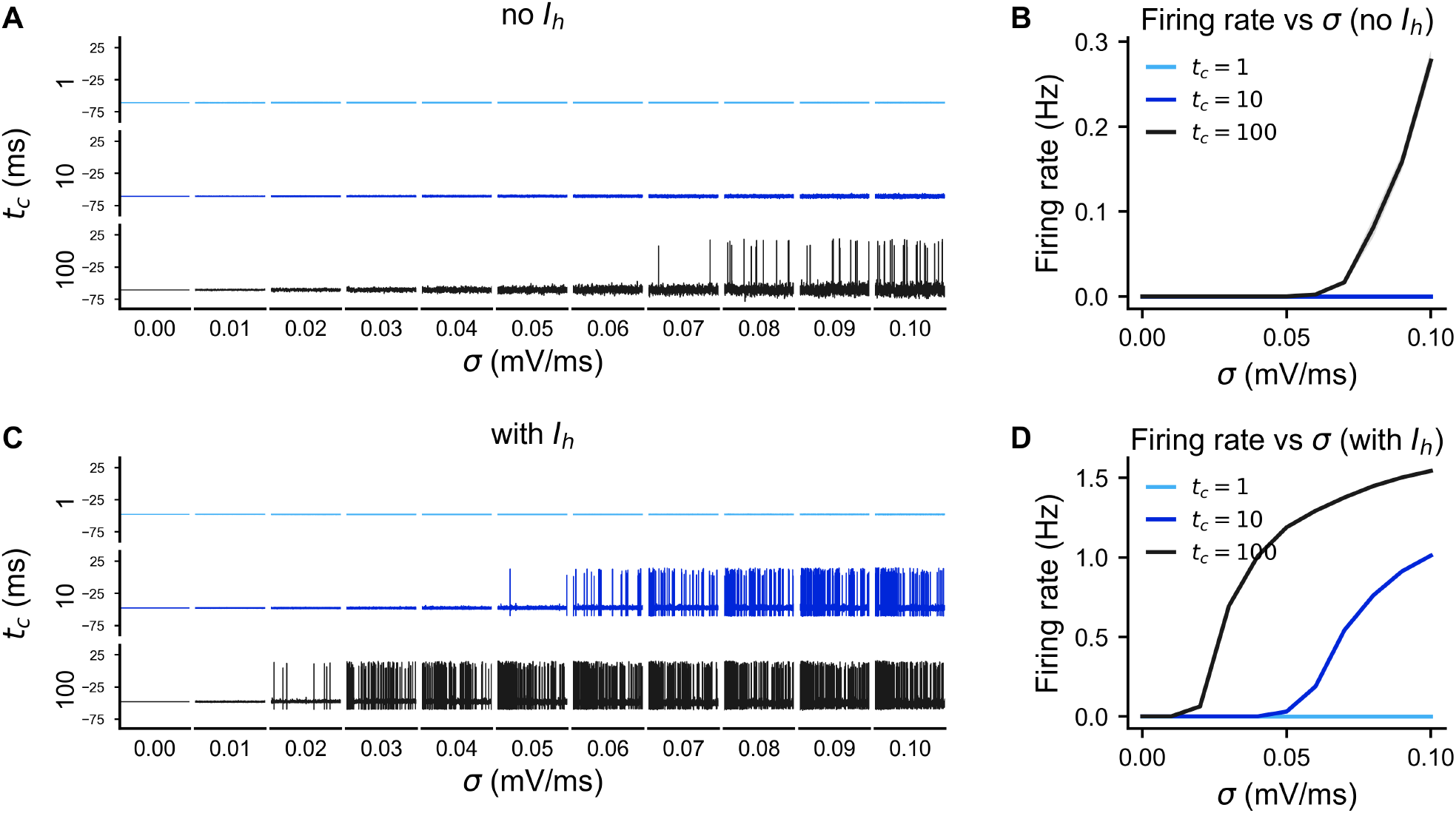
Contribution of the hyperpolarization-activated current (*I*_*h*_) to noise-driven spontaneous firing in spPVINs. (A) Representative simulations of the two-compartment Hodgkin–Huxley (HH) model in the absence of *I*_*h*_ obtained across a range of noise amplitudes (*σ* = 0.01–0.1 mV ms^*−*1*/*2^, in increments of 0.01) and three temporal correlation timescales (*t*_*c*_ = 1, 10, 100 ms). (B) Firing-rate profiles corresponding to the simulated traces shown in A, with the blue and cyan traces for *t*_*c*_ = 1 and 10 ms, respectively, overlaid on top of each other. (C) The same as in A, except that *I*_*h*_ is included in the HH model. (D) Firing-rate profiles corresponding to the simulated traces shown in C.

These results show that the temporal structure of membrane fluctuations, in addition to their amplitude, is an important determinant of spontaneous firing patterns, and that *I*_*h*_ substantially increases the sensitivity of PVINs to these fluctuations. They also suggest that differences in *I*_*h*_ expression and/or in the magnitude and temporal structure of synaptic fluctuations could contribute to distinguishing between quiescent PVINs and spPVINs.

### Role of *I*_*m*_ in shaping spPVIN activity

With the noise-driven bursting mechanism established, we next asked which intrinsic components are able to modulate the spontaneous firing activity of spPVINs. Two candidate mechanisms were considered: the outward M-type K^+^ current (*I*_*m*_) and intracellular Ca^2+^ buffering (*B*_tot_), known to modulate the mAHP via the small-conductance K^+^ current (*I*_*SK*_). On the basis of the mAHP analysis (Figure 3C), increasing *B*_tot_ reduces mAHP area, which should in principle lower the barrier to subsequent spiking. Increasing 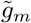, on the other hand, is expected to suppress firing by stabilizing the subthreshold membrane potential.

Parameter sweeps under stochastic drive (*I*_app_ = 0 pA, *t*_*c*_ = 100 ms, *σ* = 0.05 mV ms^*−*1*/*2^) confirm that 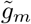 dominates over *B*_tot_ across all metrics examined (Figure 6). More specifically, firing rate decreases with 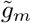 and increases with *B*_tot_ (Figure 6A), an expected outcome in view of the fact that *I*_*m*_ is an outward current [34, 35], whereas a larger *B*_tot_ attenuates the activation of the outward current *I*_*SK*_. However, when 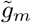 is large enough (closer to 10 nS), the HH model becomes silent, rendering variations in *B*_tot_ ineffective at restoring or modulating firing.

**Fig. 6.**
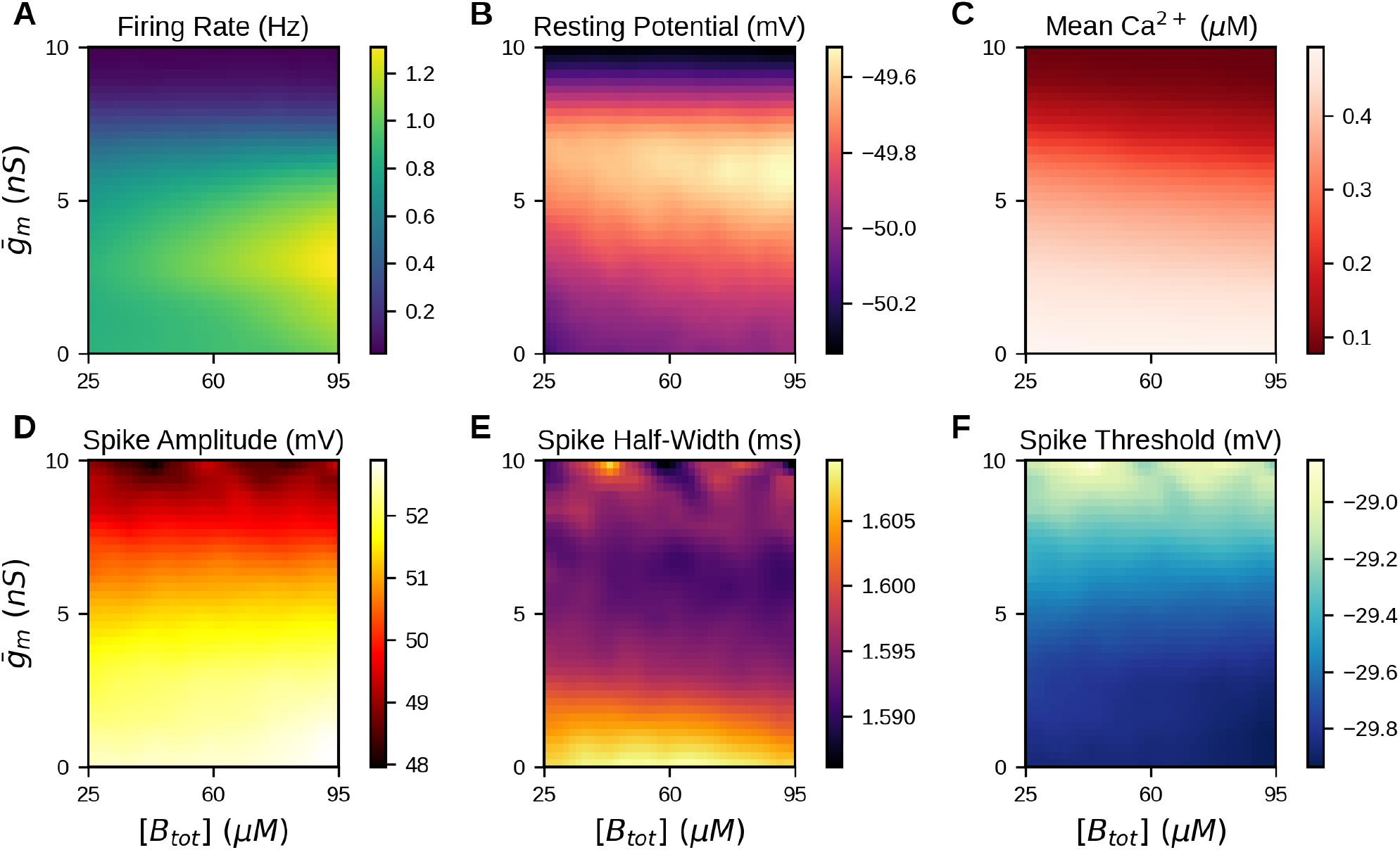
The M-type K+ current (Im) in the Hodgkin–Huxley (HH) model exerts negative feedback on spPVIN excitability. Heatmaps showing the joint effects of the Im maximum conductance 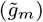 and the intracellular buffer concentration (Btot) on six electrophysiological features extracted from the HH model, including (A) firing rate, (B) resting membrane potential, (C) mean intracellular Ca2+ concentration, (D) spike amplitude, (E) spike half-width, and (F) spike threshold.

This dominant influence of 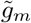 extends beyond firing rate to several other metrics of intrinsic excitability. Indeed, the resting membrane potential is strongly modulated by 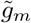, reaching its maximum at intermediate 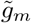 values, with little contribution from *B*_tot_ (Figure 6B). Mean intracellular Ca^2+^ concentration likewise depends more strongly on 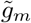 than on *B*_tot_ (Figure 6C), with a reduction in 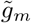 raising firing frequency and thereby increasing cumulative Ca^2+^ influx, an effect that outweighs passive buffering. The same conclusion holds for spike amplitude (Figure 6D) and threshold (Figure 6F), where reduced 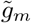 consistently shifts the neuron toward higher excitability, whereas the effect of 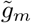 on spike half-width is biphasic (Figure 6E). Ca^2+^ buffering exerts its influence mainly through the Ca^2+^-activated conductances that shape the AHP, and therefore acts on interspike intervals more than on spike initiation itself. Across the range of *B*_tot_ explored here, this translates into a weaker effect on firing rate than that of 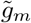, although buffering is not without effect on excitability: by attenuating the mAHP it can modestly increase firing rate, particularly at low 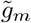. Based on this analysis, we conclude that 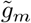 sets the excitability landscape that determines whether stochastic fluctuations can drive the neuron to threshold, while *B*_tot_ plays a secondary modulatory role.

### *I*_*h*_ promotes spontaneous firing through subthreshold depolarization

It is well known that *I*_*h*_ provides an inward depolarizing drive to neurons, bringing their membrane potential closer to firing threshold [36, 37], and promotes spontaneous firing. We therefore next examined how variations in 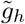 and *B*_tot_ together influence spontaneous firing and other measures of intrinsic excitability.

Parameter sweeps under stochastic drive (*I*_app_ = 0 pA, *t*_*c*_ = 100 ms, *σ* = 0.05 mV ms^*−*1*/*2^; Figure 7) demonstrated that 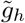exerts a strong influence on spontaneous activity. Specifically, increasing 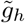 reliably increases the spontaneous firing rate and depolarizes the resting membrane potential (Figure 7A and B, respectively), causing the mean intracellular Ca^2+^ concentration to increase (Figure 7C), likely through greater cumulative Ca^2+^ influx associated with the higher firing rate. The resulting increase in intracellular Ca^2+^ in turn promotes *I*_*SK*_ activation, which may contribute to the reduction in spike amplitude (Figure 7D) and the increase in spike threshold (Figure 7F). Spike half-width, on the other hand, increases with 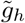 (Figure 7E), indicating broader spikes and altered repolarization kinetics. Finally, as observed previously, aside from a moderate effect of increasing *B*_tot_ on spike amplitude, Ca^2+^ buffering appears to contribute minimally to the other metrics examined.

**Fig. 7.**
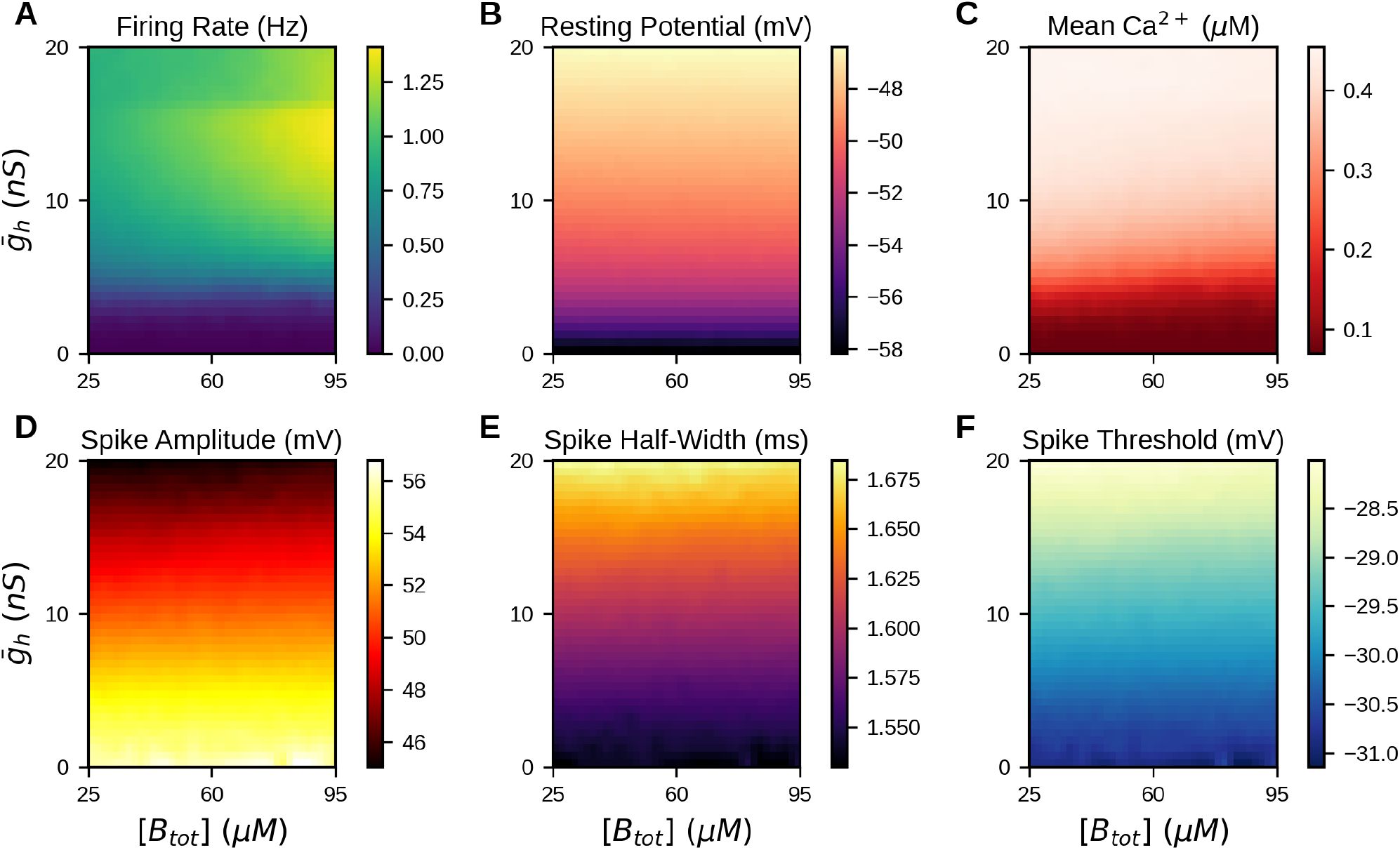
The hyperpolarization-activated current (*I*_*h*_) in the Hodgkin–Huxley (HH) model dictates spPVIN excitability. Heatmaps showing the joint effects of the *I*_*h*_ maximum conductance 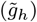 and the intracellular buffer concentration (*B*_tot_) on six electrophysiological features extracted from the HH model, including (A) firing rate, (B) resting membrane potential, (C) mean intracellular Ca^2+^ concentration, (D) spike amplitude, (E) spike half-width, and (F) spike threshold. Across these features, 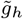 also appears dominant over *B*_tot_ in setting neuronal excitability.

Taken together, these results lead us to conclude that *I*_*h*_ plays a key role in promoting spontaneous activity, and interacts closely with *I*_*SK*_ to modulate spike features.

### Ca^2+^-driven modulation of bursting dynamics

Given that the cross-talk between *I*_*h*_ and *I*_*SK*_ appears to play a key role in shaping the excitability properties of the spPVIN HH model through Ca^2+^, we next investigated how Ca^2+^ regulates bursting in spPVINs. To do so, we first validated that [Ca^2+^]_*i*_ is a slow variable, by checking that it exhibits the typical sawtooth profile of slow variables during burst cycles (Figure 8A), and then performed a slow-fast analysis, treating [Ca^2+^]_*i*_ as a parameter to determine how it modulates the bifurcation structure of the fast subsystem [12, 41, 42]; such analysis provides a dynamical framework for understanding how the switching between active and silent phases emerges from the passage of slow variables through bifurcation points of the fast subsystem.

**Fig. 8.**
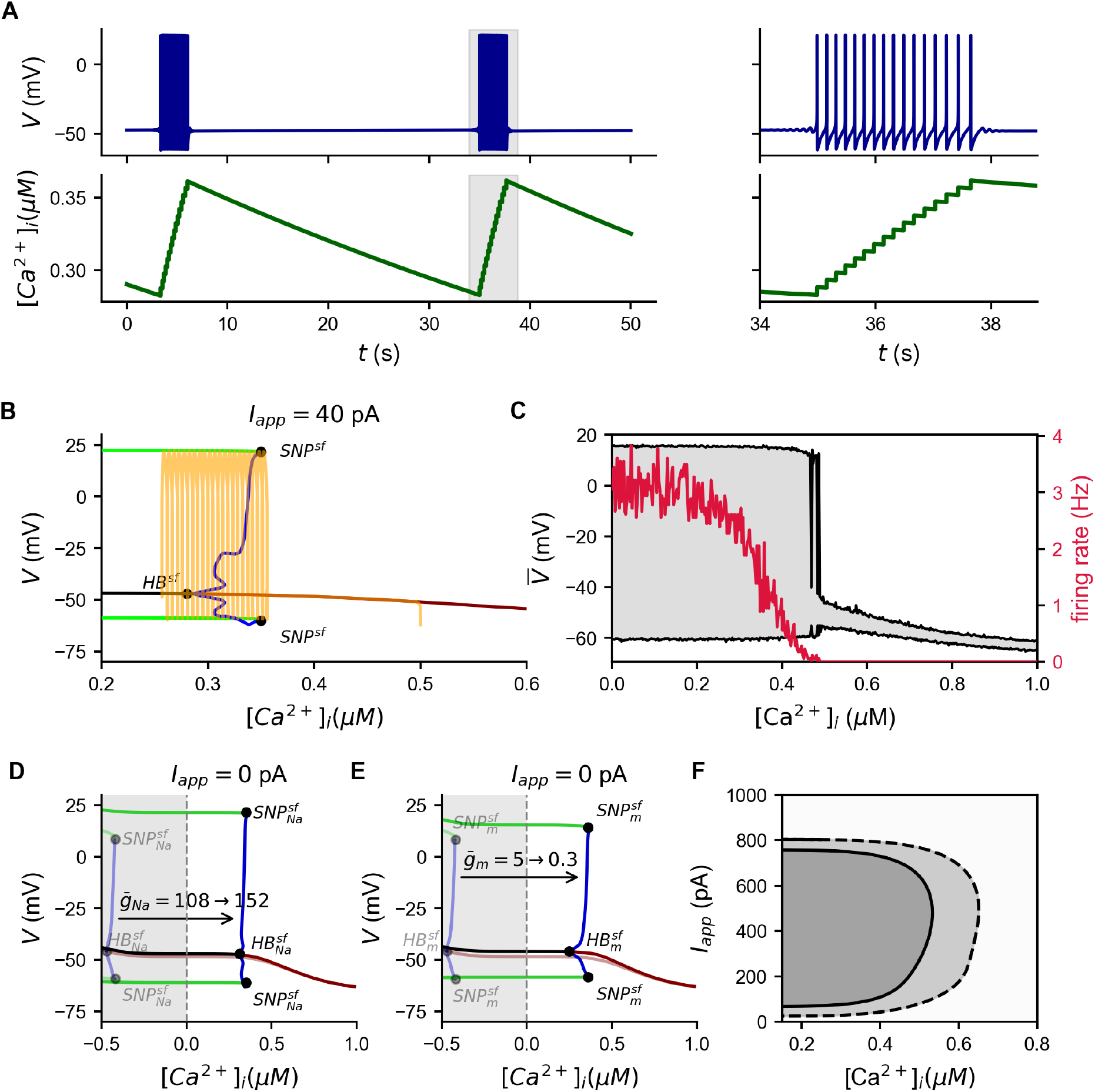
Ca^2+^-dependent elliptic bursting in the Hodgkin–Huxley (HH) model of spPVINs: a slow-fast analysis. (A) Time course of the somatic membrane potential (*V*) of the deterministic spPVIN HH model, exhibiting bursting activity (top left), and intracellular Ca^2+^ concentration ([Ca^2+^]_*i*_), showing a slow sawtooth profile (bottom left), both obtained when *I*_*app*_ = 40 pA. A zoomed-in view of a single burst cycle in *V* (top right) and the corresponding accumulation of [Ca^2+^]_*i*_ (bottom right). The burst cycle shown in the right panels is highlighted by the gray box in the left panels. (B) Slow-fast analysis of the deterministic HH model obtained by plotting the bifurcation diagram of *V* with respect to the slow variable [Ca^2+^]_*i*_, treated as a parameter, with *I*_app_ = 40 pA. Red (black) lines: branches of stable (unstable) equilibria; green (blue) lines: envelopes of stable (unstable) limit cycles. Yellow line: bursting orbit of the full system superimposed on the bifurcation diagram. HB^*sf*^ : Hopf bifurcation; SNP^*sf*^ : saddle-node bifurcation of periodic orbits. (C) The average minimum-to-maximum range of somatic membrane voltage (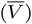) in the stochastic HH model as a function of the slow variable [Ca^2+^]_*i*_, treated as a parameter, with *I*_app_ = 0 pA (gray region), and the corresponding average firing rate (red line). (D) Bifurcation diagrams of *V* with respect to the slow variable [Ca^2+^]_*i*_, treated as a parameter, with *I*_app_ = 0 pA, for 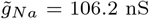 and 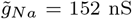. Color coding of each line is identical to that used in panel B. Gray region corresponds to the unphysiological range. 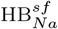: Hopf bifurcations; 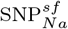 : saddle-node bifurcations of periodic orbits. (E) The same as in D, except that the two bifurcation diagrams are plotted for 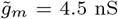 and 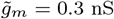. 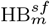: Hopf bifurcations; 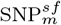 : saddle-node bifurcations of periodic orbits. (F) Two-parameter bifurcation diagram of the HH model with respect to ([Ca^2+^]_*i*_,*I*_*app*_). The solid line corresponds to Hopf bifurcations, while the dashed line corresponds to saddle-node bifurcations of periodic orbits. White: quiescent regime; light gray: bistability regime; dark gray: bursting regime.

By computing the fast-subsystem bifurcation diagram as a function of [Ca^2+^]_*i*_ at *I*_app_ = 40 pA (Figure 8B), we found that, at low [Ca^2+^]_*i*_, envelopes of stable limit cycles surround a branch of unstable equilibria. As [Ca^2+^]_*i*_ increases, these stable limit-cycle envelopes terminate at saddle-node bifurcations of periodic orbits (SNP^*sf*^), from which envelopes of unstable limit cycles emerge. The latter terminate at a subcritical Hopf bifurcation (HB^*sf*^), where the branch of unstable equilibria regains stability, giving rise to a branch of stable equilibria at high [Ca^2+^]_*i*_. The subcritical nature of HB^*sf*^ creates a hysteresis with a bistable region, where stable limit cycles and stable equilibria coexist. This bistability is central to the generation of bursting behavior. Indeed, when a burst cycle of the full system (i.e., when [Ca^2+^]_*i*_ is treated as a dynamic variable) is overlaid on the fast-subsystem bifurcation diagram, the trajectory traces a closed orbit through the stable components of this structure. Specifically, at low [Ca^2+^]_*i*_, the trajectory oscillates between envelopes of stable limit cycles, generating the active phase with periodic spiking, while simultaneously moving toward higher [Ca^2+^]_*i*_ as Ca^2+^ accumulates during spiking. Upon reaching SNP^*sf*^, the stable limit cycles disappear, terminating spiking and causing the trajectory to transition in a damped manner toward the stable equilibrium branch, thereby initiating the silent phase. During this phase, [Ca^2+^]_*i*_ decreases through Ca^2+^ efflux, allowing the trajectory to pass through HB^*sf*^ and to be gradually repelled from the unstable equilibrium branch toward the stable limit-cycle envelopes, re-initiating the next active phase. This slow passage through the Hopf bifurcation and the associated hysteresis between the active and silent states are characteristic of elliptic bursting, exhibiting mixed-mode small- and large-amplitude oscillations [12, 43].

To determine how the deterministic bifurcation structure shapes the stochastic dynamics, we incorporated OU noise into the fast subsystem, treating [Ca^2+^]_*i*_ as a parameter, and computed the average minimum-to-maximum range of the somatic membrane potential (*V*) over 10 realizations for each value of [Ca^2+^]_*i*_. The resulting diagram (Figure 8C) revealed a structure closely resembling that of the deterministic fast-subsystem bifurcation diagram (Figure 8B), with two distinct voltage envelopes. At higher [Ca^2+^]_*i*_, a narrow voltage envelope emerges, corresponding to quiescent dynamics with near-zero firing rates, whereas at lower [Ca^2+^]_*i*_, a broad voltage envelope corresponds to active spiking with high firing rates. Near the transition between these regimes, the voltage envelope broadens while the firing rate assumes intermediate values, indicating a region in which the system intermittently transitions between spiking and quiescent states. Thus, despite the stochastic nature of the dynamics, the voltage envelope and firing-rate profile recapitulate the key features of the deterministic slow-fast structure, suggesting that OU noise can drive transitions between the active and silent states along a Ca^2+^-dependent trajectory analogous to that underlying deterministic bursting.

The preceding analysis shows that bursting requires HB^*sf*^ and the associated envelopes of limit cycles of the fast subsystem to lie within the physiologically accessible region [Ca^2+^]_*i*_ *≥* 0. At *I*_app_ = 0 pA, this condition can be achieved by appropriately shifting the fast-subsystem bifurcation structure through changes in intrinsic conductances. Specifically, increasing the maximum *I*_*Na*_ conductance 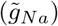 from its default value of 106.2 to 152 nS moves HB^*sf*^ into the [Ca^2+^]_*i*_ *≥* 0 regime (Figure 8D), producing a bifurcation diagram identical in structure to that obtained when *I*_app_ = 40 pA (Figure 8B), thereby enabling intrinsic bursting. Conversely, reducing the maximum *I*_*m*_ conductance 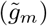 from its default value of 4.5 to 0.3 nS produces a similar shift of the bifurcation structure into the physiological region (Figure 8E), also permitting intrinsic bursting, consistent with the dominant role of *I*_*m*_ in setting the excitability landscape identified above. These results thus provide a mechanistic interpretation of the stochastic dynamics observed above (Figure 8C), namely that although *I*_app_ = 0 pA, OU noise can effectively promote transitions into the bursting regime (through channel noise) by driving the system into the same Ca^2+^-dependent bifurcation structure that is deterministically accessible through increased 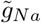 or decreased 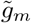. Interestingly, plotting the two-parameter bifurcation diagram of the fast subsystem with respect to both *I*_app_ and [Ca^2+^]_*i*_ further showed that this model possesses a bistable regime (Figure 8F), suggesting that OU noise may move the spPVIN HH model repetitively between bursting and quiescence.

### PVIN excitability governs inhibitory gating in the dorsal horn circuit

We have so far established how intrinsic conductances and stochastic fluctuations shape spPVIN excitability at the single-cell level. We next examined how these intrinsic properties shape spPVIN function within the dorsal horn inhibitory circuit (Figure 9A, top). Having established that *I*_*h*_ plays a critical role in regulating spPVIN excitability at the single-cell level (Figures 4, 5, and 8), we asked whether its absence alters or abolishes spPVIN-mediated feed-forward inhibition of PKC*γ* neurons within the mechanosensory circuit (Figure 9A, bottom). Specifically, we examined how the reduced excitability of spPVINs in the absence of *I*_*h*_ affects the transmission of A*β* afferent input through this circuit.

**Fig. 9.**
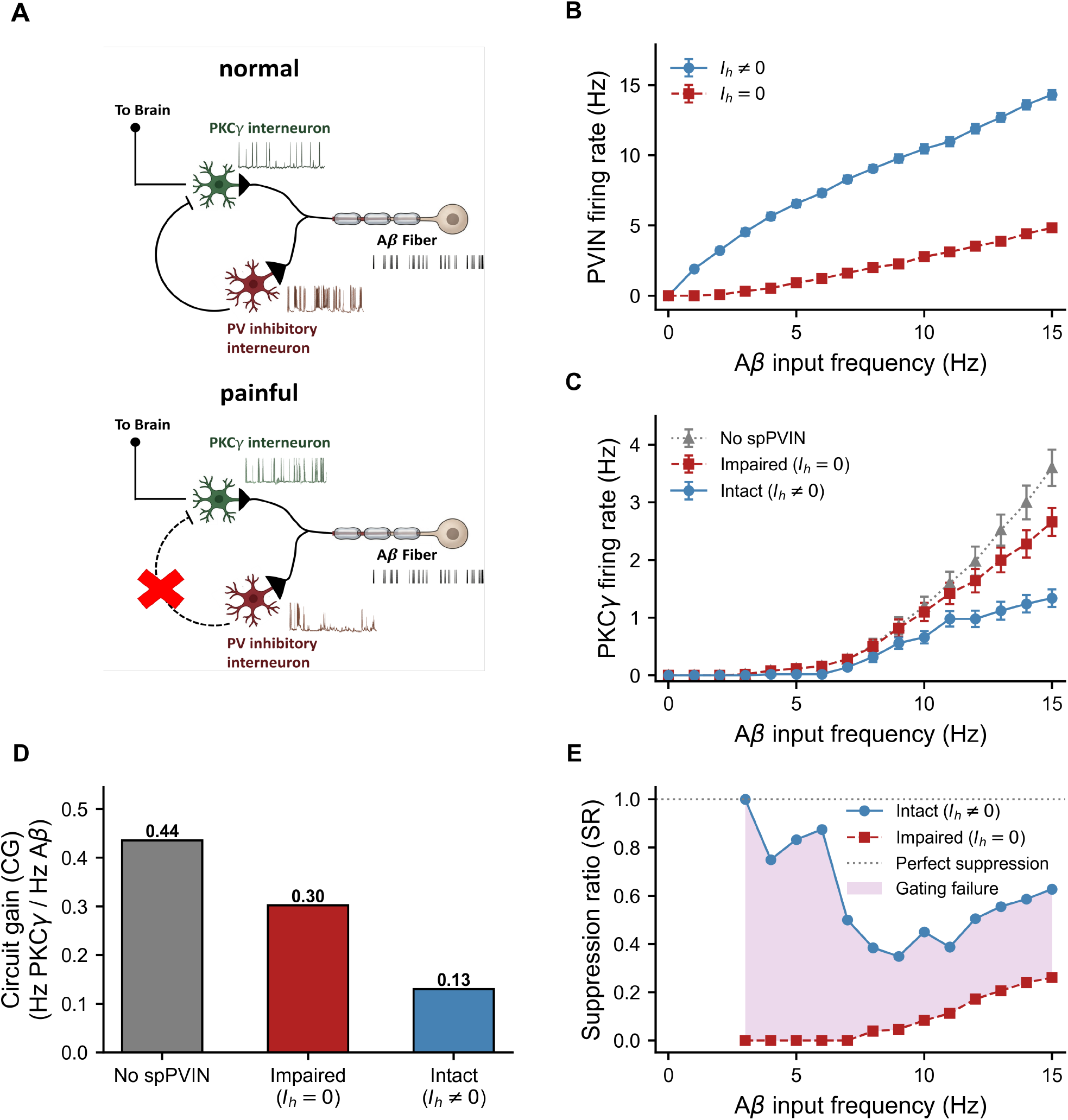
The hyperpolarization-activated current (*I*_*h*_) in the Hodgkin–Huxley (HH) model of spPVINs plays a key 1 role in enabling these neurons to suppress PKC*γ* activity. (A) Schematic of the dorsal horn mechanosensory circuit in its intact form (top; innocuous), as well as in the absence of PVIN feed-forward inhibition onto PKC*γ* (bottom; painful). A detailed description of this circuit was provided above (Figure 2A). (B,C) Average firing rates of spPVIN (B) and PKC*γ* (C) across increasing A*β* input frequencies (0–15 Hz). Lines are color-coded according to the legend. (D,E) Quantification of PKC*γ* neural responses using two measures: circuit gain computed over the 0–15 Hz range of A*β* fiber input (D), and suppression ratio (E) under three conditions: feed-forward inhibition from spPVINs expressing or lacking *I*_*h*_, and the absence of feed-forward inhibition.

To address this question, we simulated the dorsal horn mechanosensory circuit under two conditions: with *I*_*h*_ included in the spPVIN HH model and with *I*_*h*_ removed. We then quantified the average firing rate of spPVINs across 10 independent realizations, each lasting 2.5 s, in response to A*β* input frequencies ranging from 0 to 15 Hz (Figure 9B). Increasing A*β* input frequency produced a progressively greater increase in the firing rate of spPVINs expressing *I*_*h*_ compared with those lacking *I*_*h*_ (Figure 9B), demonstrating that the presence of *I*_*h*_ enhances spPVIN responsiveness to afferent input. These two conditions thus provide representative intact and impaired states of spPVIN excitability, respectively.

We next examined how this reduction in spPVIN responsiveness in the absence of *I*_*h*_ propagates through the inhibitory circuit to affect PKC*γ* neuron activity. When *I*_*h*_ was present in the spPVIN HH model, PKC*γ* firing rate and circuit gain (*CG*) remained strongly suppressed across A*β* input frequencies (Figure 9C and D), consistent with effective feed-forward inhibitory gating. In contrast, removing *I*_*h*_ from the spPVIN model weakened this inhibition and increased both PKC*γ* firing rate and *CG*, with the increase in firing rate being most pronounced at intermediate-to-high A*β* input frequencies. Notably, when compared with the condition in which feed-forward inhibition was completely removed, the PKC*γ* firing rate and *CG* in the absence of spPVIN *I*_*h*_ approached the corresponding activity levels observed without spPVIN-mediated inhibition, whereas both measures remained substantially suppressed when *I*_*h*_ was present (Figure 9C and D). We further quantified this loss of inhibition using the suppression ratio (*SR*), which showed a pronounced decrease at intermediate-to-high A*β* input frequencies when *I*_*h*_ was removed from the spPVIN HH model (Figure 9E). This reduction in *SR* indicates that hypoexcitable spPVINs become progressively less effective at suppressing PKC*γ* firing in response to increasing A*β* input. Taken together, these results demonstrate that the intrinsic excitability of spPVINs, governed in large part by *I*_*h*_, is a critical determinant of feed-forward inhibition in the dorsal horn circuit. Thus, reduced *I*_*h*_-dependent excitability can weaken spPVIN-mediated inhibitory gating and promote downstream PKC*γ* hyperexcitability.

## Discussion

The present study identified a set of intrinsic and stochastic mechanisms that shape spontaneous activity in PVINs and demonstrated how these cellular properties propagate to influence sensory gating within the dorsal horn. PVINs are predominantly quiescent under baseline conditions, yet a small subset exhibits spontaneous activity, including sparse firing and burst-like discharges. Similar heterogeneity in spontaneous activity has been observed among superficial dorsal horn neurons, where spontaneously active neurons can exhibit isolated spikes as well as distinct bursting patterns [44]. We referred to these spontaneously active PVINs as spPVINs. Their variable activity raises an important mechanistic question: why does only a subset of PVINs enter a spontaneously active state in the absence of external current? Our results suggest that differences in intrinsic conductances, particularly *I*_*h*_, *I*_*m*_ and *I*_*Na*_, together with stochastic synaptic fluctuations, can provide a mechanistic basis for this variability.

The two-compartment HH model developed here reproduced key electrophysiological features of spPVINs, including axonal spike initiation and the characteristic kink at spike onset. The importance of the axon initial segment (AIS) in determining spike threshold and action-potential initiation is well established [45, 46]. Incorporating a distinct AIS compartment therefore provided an important spatial constraint on the model and allowed the intrinsic mechanisms governing somatic excitability to be examined without neglecting the specialized geometry of spike initiation. More importantly, our results indicate that spontaneous activity does not simply emerge from membrane noise acting on an otherwise passive or uniformly excitable membrane. Rather, 8 stochastic fluctuations interact with a structured set of intrinsic conductances to determine whether the neuron 9 remains quiescent, generates isolated spikes, or enters a bursting regime.

In particular, adding OU noise directly to the membrane-voltage equation allowed us to examine how temporally correlated fluctuations interact with intrinsic excitability. We found that the temporal persistence of the fluctuations is an important determinant of firing behavior, with short-lived fluctuations preferentially producing isolated action potentials, consistent with a shift toward type II, resonator-like excitability under *in vivo*-like conditions [47], whereas larger or more persistent fluctuations increasingly promote burst firing. This result is consistent with recent work on electrosensory lateral-line-lobe (ELL) pyramidal cells showing that heterogeneous firing patterns emerge from the interaction between intrinsic properties and stochastic synaptic inputs [18, 19]. The similarity between these observations and our results suggests that stochastic input may be an important determinant of whether an intrinsically capable neuron remains quiescent, fires sparsely, or enters a bursting state. Importantly, our model adds a mechanistic interpretation of this phenomenon by showing that temporally correlated fluctuations can hold the membrane potential near threshold for sufficiently long periods to recruit slow intrinsic currents, thereby facilitating the transition from isolated spiking to bursting.

The stochastic mechanism may also help explain why only a small subset of PVINs exhibits spontaneous activity under baseline conditions. One possibility is that these spPVINs possess greater functional expression of HCN channels and consequently stronger *I*_*h*_. HCN channels are expressed in PV interneurons and can influence their functional output [5, 48], while *I*_*h*_ provides a depolarizing current that is recruited following membrane hyperpolarization and can promote neuronal excitability [36, 37]. Our simulations support this possibility by showing that increasing 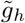 depolarizes the resting membrane potential, reduces the distance to spike threshold, and substantially lowers the stochastic drive required to elicit spontaneous firing. Alternatively, spPVINs may receive stronger or more persistent synaptic input, thereby experiencing greater fluctuations in membrane potential that facilitate spontaneous activity. PV interneurons in other circuits can receive substantial convergent excitatory input and exhibit extensive synaptic connectivity [49]. Although these possibilities remain to be tested directly in dorsal horn PVINs, they suggest that both intrinsic *I*_*h*_-dependent excitability and the strength or temporal structure of synaptic input may contribute to the emergence of spontaneous activity in spPVINs.

Our parameter and bifurcation analyses identified *I*_*m*_, *I*_*h*_, and *I*_*Na*_ as key determinants of how stochastic fluctuations are translated into firing. *I*_*m*_ acts as an outward brake, whereas *I*_*h*_ provides a depolarizing drive, with their balance establishing the excitability landscape. At low 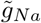, the model exhibits a stable equilibrium with type III excitability, allowing perturbations to elicit isolated spikes without sustained repetitive firing [50]. Increasing 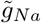 expands the regime in which repetitive firing and bursting are possible, whereas increasing 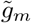 promotes quiescence. Thus, OU noise can transiently move the model out of the quiescent regime and into the bursting regime, consistent with previous work showing that channel noise can interact with intrinsic dynamics to promote transitions between spiking and bursting [51]. These findings suggest that stochastic fluctuations arising from synaptic input and/or channel noise may enable PVINs to switch between quiescent, isolated-spiking, and bursting states without requiring sustained external depolarization.

Slow-fast analysis provided a mechanistic explanation for the emergence of elliptic bursting in the model. Because the intracellular Ca^2+^ concentration ([Ca^2+^]_*i*_) evolves on a much slower timescale than the other variables in the HH model, it can be treated as a parameter when analyzing the remaining voltage and gating dynamics as a fast subsystem, an approach commonly used to characterize neuronal bursting [43, 52] and previously applied to modeling PVINs [8]. Ca^2+^ accumulation then moves the fast subsystem between spiking and quiescent regimes, generating bursting through the interaction of fast voltage dynamics and slow Ca^2+^ evolution. The accessibility of this bursting regime is strongly regulated by 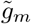 and 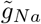, which determine the location of the Hopf and saddle-node bifurcations of periodic orbits.

Together, these analyses identify a hierarchy of intrinsic mechanisms controlling spPVIN activity. *I*_*m*_ and *I*_*h*_ establish the baseline excitability landscape, whereas *I*_*Na*_ and the slow Ca^2+^ dynamics determine whether that landscape supports repetitive spiking and bursting. *B*_tot_, in contrast, primarily modulates the consequences of firing by shaping the mAHP through *I*_*SK*_ and thereby refining interspike intervals. This distinction is important because it suggests that Ca^2+^ buffering does not primarily determine whether a PVIN fires, but rather modifies the temporal structure of firing once the neuron has entered an active regime. The model therefore separates the mechanisms controlling entry into spontaneous activity from those controlling the temporal organization of that activity.

The functional importance of these intrinsic mechanisms becomes apparent at the circuit level. PVINs are central components of the dorsal horn inhibitory network and are thought to regulate the transmission of innocuous tactile information toward ascending nociceptive pathways [29, 31, 53]. Under baseline conditions, activation of low-threshold A*β* afferents reliably recruits PVINs, which in turn impose strong feed-forward inhibition onto PKC*γ* neurons. This inhibitory gate prevents innocuous tactile input from being transmitted 6 through nociceptive pathways [29, 31]. Our circuit simulations show that the intrinsic excitability of spPVINs 7 is an important determinant of the strength of this gate. When *I*_*h*_ is present, increasing A*β* input progressively 8 promotes spPVIN firing, producing strong inhibition of PKC*γ* neurons. In contrast, removal of *I*_*h*_ reduces 9 spPVIN responsiveness, weakens feed-forward inhibition, and allows A*β* input to drive PKC*γ* activity.

This provides an important link between the single-cell and circuit-level analyses. Unlike the approach of Ma et al. [8], which modeled impaired inhibition primarily through changes in *B*_tot_ and consequent spike-2 frequency adaptation, we focused here on the intrinsic excitability of PVINs, specifically the contribution of *I*_*h*_, 3 as a mechanism regulating feed-forward inhibition. Based on our single-cell analysis, *I*_*h*_ depolarizes the resting 4 membrane potential and reduces the distance to spike threshold [36, 37]. Consequently, its removal decreases the 5 ability of PVINs to respond reliably to A*β* input. The resulting reduction in PVIN responsiveness is sufficient 6 to weaken inhibitory gating and disinhibit PKC*γ* neurons. Indeed, when *I*_*h*_ is removed, both PKC*γ* firing and 7 circuit gain approach the responses observed when PVIN-mediated feed-forward inhibition is completely absent. 8 This demonstrates that a change in a single intrinsic conductance can propagate from the membrane level to 9 the circuit level and substantially alter sensory processing.

The circuit-level consequences are particularly relevant to neuropathic pain. PVIN-mediated inhibition normally acts as a gate that prevents low-threshold tactile input from accessing nociceptive pathways. Following 2 nerve injury, impaired PVIN-mediated inhibition has been implicated in the emergence of mechanical allody-3 nia [1, 53]. Our results suggest that reduced intrinsic excitability of PVINs represents one mechanism through 4 which this inhibitory gate could become compromised. If changes in channel expression or function shift PVINs 5 from a reliably responsive state toward sparse firing or quiescence, A*β* input would be less effectively suppressed, 6 increasing the likelihood that tactile information recruits downstream nociceptive circuitry. Thus, a decrease in 7 spPVIN excitability provides a potential cellular mechanism linking altered intrinsic membrane properties to 8 the dorsal horn disinhibition and hyperexcitability associated with neuropathic pain [1, 8].

More broadly, the present study illustrates the value of combining biophysical modeling, stochastic dynamics, and bifurcation analysis to connect cellular mechanisms with circuit function. Rather than treating spontaneous 1 firing as a phenomenological property, the model identified how a small set of conductances determines the 2 accessibility of qualitatively distinct dynamical regimes and how stochastic fluctuations can move the neuron 3 between these regimes. The two-compartment architecture further linked these intrinsic dynamics to the spatial 4 organization of spike initiation, while the slow-fast analysis provided a mechanistic explanation for the emergence 5 of bursting. Finally, embedding the model within the dorsal horn circuit demonstrated how changes in single-6 cell excitability could propagate to alter sensory gating. This multiscale framework provides a basis for future 7 studies incorporating activity-dependent channel plasticity, injury-induced changes in conductance densities, 8 heterogeneous synaptic connectivity, and state-dependent neuromodulation to determine how alterations in 9 PVIN intrinsic properties evolve into persistent circuit dysfunction during chronic pain.

## Declarations

### Data availability

All data supporting the findings of this study are available from the corresponding author upon reasonable 6 request.

### Code availability

All Python code and XPPAUT files used to generate and analyze the model and circuit simulations are publicly 0 available at https://github.com/rfritzdj/spPVIN.

### Author contributions

4 R.F.D.J., R.S.-N. and A.K. designed the study. R.F.D.J. developed the model and performed the simulations 5 and analyses. C.T. and H.Q. performed the electrophysiological recordings. E.C., A.Kr., R.S.-N. and A.K. 6 supervised the work. R.F.D.J. and A.K. wrote the manuscript with input from all authors.

## Acknowledgements

We thank the staff of the Comparative Medicine and Animal Resources Centre at McGill University for their management of the mouse colony.

