## Supplementary Material for "Intrinsic Ionic Mechanisms Underlying Burst Firing in Spontaneously Active Dorsal Horn Parvalbumin Interneurons"

#### Appendix S1 PKC$\gamma$ model: membrane potential equation

The PKC$\gamma$ neuron was modeled as a single isopotential somatic compartment, adapted from the superficial dorsal horn model of Medlock et al. [28]. The soma is treated as a cylinder of length $L$ and diameter $D_{\text{soma}}$ (Table 2 of the main text), giving a surface area $A_\gamma = \pi L D_{\text{soma}}$.

The membrane potential $V_\gamma$ satisfies

$$C_{m,\gamma} \frac{dV_\gamma}{dt} = -I_{Na1} - I_{Na2} - I_K - I_A - I_{KDR} - I_{\text{leak}} - \frac{\chi}{A_\gamma}\left(I_{exc} + I_{inh}\right), \tag{S1}$$

where $I_{Na1}$ and $I_{Na2}$ are the two HH-type $Na^+$ currents [11, 54], $I_K$, $I_A$ and $I_{KDR}$ are the HH-type [11], A-type [55] and delayed-rectifier $K^+$ [56] currents, respectively, $I_{\text{leak}}$ is a nonspecific leak current, $I_{exc}$ is the excitatory synaptic current (AMPA and NMDA receptor-mediated, driven by A$\beta$ afferents), and $I_{inh}$ is the inhibitory synaptic current (glycine and $GABA_A$ receptor-mediated, driven by spPVIN spikes). The two synaptic currents are computed as absolute currents (in pA; Section S3), whereas the intrinsic currents are expressed as densities (in $\mu$A/cm$^2$); the factor $\chi = 100\ \mu\text{A}\,\text{cm}^{-2}\,\text{pA}^{-1}\mu\text{m}^2$ converts the former to the latter, giving $\chi/A_\gamma \simeq 0.0796\ \mu\text{A}\,\text{cm}^{-2}\,\text{pA}^{-1}$ for the geometry used here. No external current was injected into the PKC$\gamma$ model in any of the circuit simulations reported.

Each voltage-gated current takes the canonical conductance-based form,

$$I_{Na1} = \tilde{g}_{Na1} \cdot m_{Na1}^3 h_{Na1} \cdot (V_\gamma - E_{Na}), \tag{S2}$$

$$I_{Na2} = \tilde{g}_{Na2} \cdot m_{Na2}^3 h_{Na2} \cdot (V_\gamma - E_{Na}), \tag{S3}$$

$$I_K = \tilde{g}_K \cdot n_K^4 \cdot (V_\gamma - E_K), \tag{S4}$$

$$I_{KDR} = \tilde{g}_{KDR} \cdot n_{KDR}^4 h_{KDR} \cdot (V_\gamma - E_K), \tag{S5}$$

$$I_A = \tilde{g}_A \cdot n_A l_A \cdot (V_\gamma - E_K), \tag{S6}$$

$$I_{\text{leak}} = \tilde{g}_{\text{leak}} \cdot (V_\gamma - E_{\text{leak}}), \tag{S7}$$

where the maximal conductances $\tilde{g}_j$ and reversal potentials $E_j$ are listed in Table 2 of the main text. All gating variables $x \in \{m_{Na1}, h_{Na1}, m_{Na2}, h_{Na2}, n_K, n_A, l_A, n_{KDR}, h_{KDR}\}$ follow first-order kinetics,

$$\frac{dx}{dt} = \frac{x_\infty - x}{\tau_x}, \tag{S8}$$

with $x_\infty$ and $\tau_x$ specified current-by-current in Section S2. Unless stated otherwise there, steady states and time constants are obtained from the forward and backward rates as

$$x_\infty = \frac{\alpha_x}{\alpha_x + \beta_x}, \qquad \tau_x = \frac{1}{(\alpha_x + \beta_x)\, t_{adj}}, \tag{S9}$$

where $t_{adj}$ is a temperature-scaling factor whose value differs between currents (Section S2). Throughout, $V \equiv V_\gamma$ is in mV, rate constants are in ms$^{-1}$, time constants are in ms, and all simulations used $T = 23$°C. Gating variables were initialized at their steady-state values for $V_\gamma = -65$ mV.

Several of the rate expressions below have the form $c\,x/\big(1 - e^{-x/y}\big)$ or $c\,x/\big(e^{x/y} - 1\big)$ and are therefore of indeterminate form at $x = 0$. In every such case the singularity is removable, and the rate takes the limiting value $c\,y$ there.

The PKC$\gamma$ model includes no $Ca^{2+}$-activated conductances, and intracellular $Ca^{2+}$ was therefore not tracked in this compartment; $Ca^{2+}$ handling in this study was restricted to the spPVIN model, where it drives $I_{SK}$.

### Appendix S2 PKC$\gamma$ model: voltage-dependent gating kinetics

#### S2.1 HH-type 1 $Na^+$ current ($I_{Na1}$)

This current is not temperature-scaled ($t_{adj} = 1$). Activation follows Eqs. (S8)–(S9) with

$$\alpha_m = \frac{0.182\,(V+28)}{1-\exp\left(-\frac{V+28}{9}\right)}, \qquad \beta_m = \frac{0.124\,(V+28)}{\exp\left(\frac{V+28}{9}\right)-1}. \tag{S10}$$

Inactivation uses an explicitly specified steady state together with a rate-derived time constant,

$$h_\infty = \frac{1}{1+\exp\left(\frac{V+64}{9}\right)}, \qquad \tau_h = \frac{1}{\alpha_h+\beta_h}, \tag{S11}$$

$$\alpha_h = \frac{0.061\,(V+35)}{1-\exp\left(-\frac{V+35}{3}\right)} + 0.0166, \qquad \beta_h = \frac{0.0018\,(V+71)}{\exp\left(\frac{V+71}{18}\right)-1}.$$

#### S2.2 HH-type 2 $Na^+$ current ($I_{Na2}$)

This current is expressed in terms of the shifted voltage $V_2 = V+63$, with temperature scaling $t_{adj} = 3^{(T-36)/10}$:

$$\alpha_m = \frac{0.32\,(13-V_2)}{\exp\left(\frac{13-V_2}{4}\right)-1}, \qquad \beta_m = \frac{0.28\,(V_2-40)}{\exp\left(\frac{V_2-40}{5}\right)-1}, \tag{S12}$$

$$\alpha_h = 0.128\exp\left(\frac{19-V_2}{18}\right), \qquad \beta_h = \frac{4}{1+\exp\left(\frac{42-V_2}{5}\right)}.$$

Steady states and time constants follow Eq. (S9) with this $t_{adj}$.

#### S2.3 HH-type $K^+$ current ($I_K$)

With shifted voltage $V_2 = V + 50.2$ and temperature scaling $t_{adj} = 3^{(T-36)/10}$:

$$\alpha_n = \frac{0.032\,(15-V_2)}{\exp\left(\frac{15-V_2}{5}\right)-1}, \qquad \beta_n = 0.5\exp\left(\frac{10-V_2}{40}\right), \tag{S13}$$

with $n_\infty$ and $\tau_n$ given by Eq. (S9).

#### S2.4 Delayed-rectifier $K^+$ current ($I_{KDR}$)

This current is not temperature-scaled ($t_{adj} = 1$). Activation ($n$) and inactivation ($h$) both follow Eqs. (S8)–(S9) with

$$\alpha_n = \frac{0.035\,(V+15)}{1-\exp\left(-\frac{V+15}{9}\right)}, \qquad \beta_n = 0.014\exp\left(\frac{-V+12}{46}\right), \tag{S14}$$

$$\alpha_h = 0.0083\left(\frac{1}{\exp\left(\frac{V+20}{10}\right)+1}+1\right), \qquad \beta_h = \frac{0.0083}{\exp\left(\frac{-V-20}{10}\right)+1}.$$

#### S2.5 A-type $K^+$ current ($I_A$)

The A-type current uses a Borg-Graham type formulation in which the steady states and time constants are specified directly rather than through Eq. (S9). Its kinetics depend on the thermodynamic factor

$$\phi = \frac{9.648\times10^4}{8.315\,(273.16+T)} \tag{S15}$$

and the temperature-scaling factor $q_{10} = 3^{(T-30)/10}$. For activation ($n_A$),

$$\alpha_k = \exp(-0.0004\,(V+15)\,\phi)\,, \qquad n_{A,\infty} = \frac{1.2}{1+\alpha_k}, \tag{S16}$$
$$\alpha_n = \exp(-0.003\,(V+45)\,\phi)\,, \qquad \beta_n = \exp(-0.00135\,(V+45)\,\phi)\,,$$
$$\tau_{n_A} = \frac{0.1\,\beta_n}{0.04\,q_{10}\,(1+\alpha_n)}.$$

For inactivation ($l_A$),

$$\alpha_m = \exp(0.004\,(V+76)\,\phi)\,, \qquad l_{A,\infty} = \frac{1}{1+\alpha_m}, \tag{S17}$$
$$\alpha_l = \beta_l = \exp(0.002\,(V+67)\,\phi)\,, \qquad \tau_{l_A} = \frac{1.2\,\beta_l}{0.023\,q_{10}\,(1+\alpha_l)}.$$

Two features of this formulation are inherited from the original implementation and are intentional rather than typographical. First, the scaling factor of 1.2 in $n_{A,\infty}$ means that this variable is not bounded above by unity; it acts as an additional gain on $\tilde{g}_A$ at depolarized potentials. Second, $\alpha_l$ and $\beta_l$ are identical because the corresponding asymmetry parameter is set to one, so that $\tau_{l_A}$ reduces to $1.2\,\alpha_l/[0.023\,q_{10}\,(1+\alpha_l)]$.

### Appendix S3 Synaptic currents in the dorsal horn circuit

All synaptic conductances were modeled with dual-exponential kinetics. For a synapse of type $s$,

$$G_s(t) = W_s \cdot f_s \cdot \left(e^{-t/\tau_{d,s}} - e^{-t/\tau_{r,s}}\right), \tag{S18}$$

where $\tau_{r,s}$ and $\tau_{d,s}$ are the rise and decay time constants, $W_s$ is the lumped synaptic weight (in nS), and $f_s$ normalizes the transient such that $\max_t G_s(t) = W_s$. Conductance transients evoked by successive presynaptic spikes were summed linearly. Weights and kinetic parameters are listed in Table 3 of the main text.

#### S3.1 Excitatory current onto PKC$\gamma$ ($I_{exc}$)

Excitatory drive onto PKC$\gamma$ arises from A$\beta$ afferents and is mediated by AMPA and NMDA receptors, both with reversal potential $E_{exc} = 0$ mV:

$$I_{exc} = \Big[G_{AMPA}(t) + G_{NMDA}(t) \cdot Mg_{block}(V_\gamma)\Big] \cdot (V_\gamma - E_{exc})\,, \tag{S19}$$

where $G_{AMPA}$ and $G_{NMDA}$ follow Eq. (S18) and are incremented at each A$\beta$ spike time. The NMDA component includes the voltage-dependent $Mg^{2+}$ block

$$Mg_{block}(V) = \frac{1}{1 + \dfrac{[\mathrm{Mg}^{2+}]_o}{3.57}\, e^{-0.062V}}, \qquad [\mathrm{Mg}^{2+}]_o = 1 \text{ mM}, \tag{S20}$$

evaluated at the membrane potential at the start of each integration step.

#### S3.2 Inhibitory current onto PKC$\gamma$ ($I_{inh}$)

Feed-forward inhibition onto PKC$\gamma$ arises from spPVIN spikes, detected during integration whenever the spPVIN somatic membrane potential crossed $-20$ mV from below. Each detected spike triggered co-released glycine and $GABA_A$ conductance transients, both with reversal potential $E_{inh} = -70$ mV:

$$I_{inh} = \Big[G_{Gly}(t) + G_{GABA_A}(t)\Big] \cdot (V_\gamma - E_{inh})\,, \tag{S21}$$

with $G_{Gly}$ and $G_{GABA_A}$ again given by Eq. (S18). In the "no PVIN" condition, $I_{inh} \equiv 0$, which defines the disinhibited baseline used to compute the suppression ratio ($SR$) in the main text.

##### S3.3 Excitatory current onto spPVINs

The spPVIN model received only excitatory A$\beta$ input, mediated by AMPA and NMDA receptors with $E_{exc} = 0$ mV and constructed exactly as for $I_{exc}$ above, but with the spPVIN-specific weights given in Table 3 of the main text. In circuit simulations, the Ornstein–Uhlenbeck noise term $\eta$ of the single-cell spPVIN model was removed and replaced by this synaptic current.
